# A novel analgesic pathway from parvocellular oxytocin neurons to the periaqueductal gray

**DOI:** 10.1101/2022.02.23.481531

**Authors:** Mai Iwasaki, Arthur Lefevre, Ferdinand Althammer, Olga Łąpieś, Louis Hilfiger, Damien Kerspern, Meggane Melchior, Stephanie Küppers, Quirin Krablicher, Ryan Patwell, Sabine C Herpertz, Beate Ditzen, Kai Schönig, Dusan Bartsch, Javier E. Stern, Pascal Darbon, Valery Grinevich, Alexandre Charlet

**Affiliations:** Centre National de la Recherche Scientifique and University of Strasbourg, Institute of Cellular and Integrative Neuroscience, 67000 Strasbourg, France; Department of Neuropeptide Research in Psychiatry, Central Institute of Mental Health, University of Heidelberg, Mannheim 68159, Germany; Center for Neuroinflammation and Cardiometabolic Diseases, Georgia State University, Atlanta, GA, USA; Department of General Psychiatry, Center of Psychosocial Medicine, University of Heidelberg, 69115, Heidelberg, Germany; Institute of Medical Psychology, Heidelberg University Hostpital, 69115, Heidelberg, Germany; Ruprecht-Karls University Heidelberg, Heidelberg, Germany; Department of Molecular Biology, Central Institute of Mental Health, University of Heidelberg, Mannheim 68159, Germany

**Author notes:** Correspondences: Alexandre Charlet, PhD, Institute of Cellular and Integrative Neuroscience, INCI CNRS UPR3212, 8, Allée du Général Rouvillois 67000 Strasbourg, France, Valery Grinevich, MD, PhD, Department of Neuropeptide Research in Psychiatry, Central Institute of Mental Health Medical Faculty Mannheim University of Heidelberg, J5, Mannheim, 68159 Germany, mail. Equal first authors (Mai Iwasaki, Arthur Lefevre). Senior authors (Valery Grinevich, Alexandre Charlet). Institute of Human Genetics, University Hospital Heidelberg, Germany (Ferdinand Althammer).

**Keywords:** Oxytocin, oxytocin receptor, periaqueductal grey, spinal cord, inflammatory pain, neuropathic pain

## Abstract

The hypothalamic neuropeptide, oxytocin (OT), exerts prominent analgesic effects via central and peripheral action. Here we discovered a novel subset of OT neurons whose projections preferentially terminate on OT receptor (OTR)-expressing neurons in the ventrolateral periaqueductal gray (vlPAG). Using a newly generated line of transgenic rats (OTR-IRES-Cre), we determined that most of the vlPAG OTR expressing cells being targeted by OT projections are GABAergic in nature. Both optogenetically-evoked axonal OT release in the vlPAG as well as chemogenetic activation of OTR vlPAG neurons results in a long-lasting overall increase of vlPAG neuronal activity. This then leads to an indirect suppression of sensory neuron activity in the spinal cord and strong analgesia. Finally, we describe a novel OT→vlPAG→spinal cord circuit that seems critical for analgesia in the context of both inflammatory and neuropathic pain.

**Highlights:**

- We generated a new transgenic knock-in rat line (OTR-IRES-Cre)
- A distinct parvOT neuronal population projects to vlPAG but not the SON or spinal cord
- OT excites vlPAG OTR neurons which indirectly inhibit SC WDR neurons
- This novel parvOT→vlPAG→SC pathway alleviates nociception but not the affective component of pain

## Introduction

The hypothalamic neuropeptide oxytocin (OT) modulates several key neurophysiological functions, including pain modulation (Poisbeau et al., 2017). OT is produced in the hypothalamic supraoptic (SON) and paraventricular (PVN) nuclei by two major types of neurons: magnocellular (magnOT) and parvocellular (parvOT) neurons. MagnOT neurons of the SON and PVN are large cells that release OT into the bloodstream via axonal projections to the posterior pituitary. In contrast, parvOT neurons are smaller cells located exclusively in the PVN and project to the brainstem and spinal cord, but not the posterior pituitary (Althammer and Grinevich, 2017). It has been previously demonstrated that a small population of PVN parvOT neurons attenuates pain perception via two pathways: 1) through coordinated OT release into the bloodstream from magnOT neurons leading to the modulation of peripheral nociceptor activity in the dorsal root ganglion and skin and 2) by inhibiting sensory neurons in the spinal cord (Eliava et al., 2016; González-Hernández et al., 2017; Moreno-López et al., 2013).

The periaqueductal gray (PAG) plays a pivotal role in descending analgesic pathways (Melzack, 1975). Indeed, physiological suppression of pain seems to be primarily modulated by a top-down system comprised of the PAG, rostral ventromedial medulla (RVM), and dorsal horn of the spinal cord (SC) (Fields, 2000). For example, electrical stimulation of the PAG inhibits the firing rate of neurons in the dorsal horn of the spinal cord (Basbaum et al., 1977; Liebeskind et al., 1973). In addition, both OT axons and OT receptors (OTR) have been reported in the PAG of mice (Campbell et al., 2009; Nasanbuyan et al., 2018), where the administration of exogenous OT into the PAG enhances neuronal firing rates (Ogawa et al., 1992) and blocking OTRs in the PAG decreases pain threshold (Yang et al., 2011).

Altogether, these studies suggest that OT may promote analgesia through the OT-mediated activation of PAG neurons. It is therefore tempting to hypothesize two independent, yet complementary mechanisms of OT-mediated analgesia in which OT attenuates nociceptive signals at the level of peripheral nociceptors or the spinal cord (Eliava et al., 2016; Herpertz et al., 2019), and also acts within the PAG to fine-tune additional descending pain-related pathways.

However, neither the cellular circuitry nor the analgesic effects of endogenous OT signaling in the PAG have been studied. To address this gap, we first generated a new Cre knock-in rat line to label and manipulate OTR neurons in the PAG, wherein we observed both synaptic and non-synaptic contacts of OTergic axon terminals with somata and dendrites of OTR-positive PAG neurons. Next, we employed cell-type-specific viral vectors to identify a new subpopulation of parvOT neurons projecting to the ventrolateral subregion of the PAG (vlPAG). We then used in vivo electrophysiology combined with optogenetics in the vlPAG to reveal that activation of OTR neurons leads to inhibition of sensory wide dynamic range (WDR) neurons in the spinal cord (SC_WDR_) of anesthetized rats. Finally, we found that optogenetically-evoked OT release in the vlPAG produces analgesia and this effect was recapitulated by chemogenetic activation of vlPAG OTR neurons. Altogether, we identified an independent parvOT→vlPAG_OTR_→SC_WDR_ pathway that is distinct from the previously described direct parvOT→SC_WDR_ pathway (Eliava et al., 2016) and is capable of promoting analgesia in the context of both inflammatory and chronic neuropathic pain.

## Results

### vlPAG OTR-expressing neurons are GABAergic

To study OTR-expressing neurons in the vlPAG, we generated a new transgenic line of rats with Cre recombinase expression controlled by the endogenous OTR gene locus (OTR-IRES-Cre line, see Methods for details) (**Figure 1A1**, **S1A**). To label OTR neurons, we injected the PAG of OTR-IRES-Cre female rats with a rAAV carrying a Cre-dependent GFP expression cassette (rAAV1/2-pEF1a-DIO-GFP) (n = 4). We found a clustering of OTR neurons along the anteroposterior axis of the vlPAG (**Figure S1B-C**). We observed a similar pattern in males (data not shown). To further assess the specificity of Cre localization in OTR-IRES-Cre rats, we performed RNA-Scope using probes against both OTR and Cre mRNAs and found that 97.6% of Cre positive cells (n = 47) were also positive for OTR mRNAs and 90.4% of OTR positive cells (n = 42) also expressed Cre mRNAs (**Figure 1A2**).

**Figure 1.**
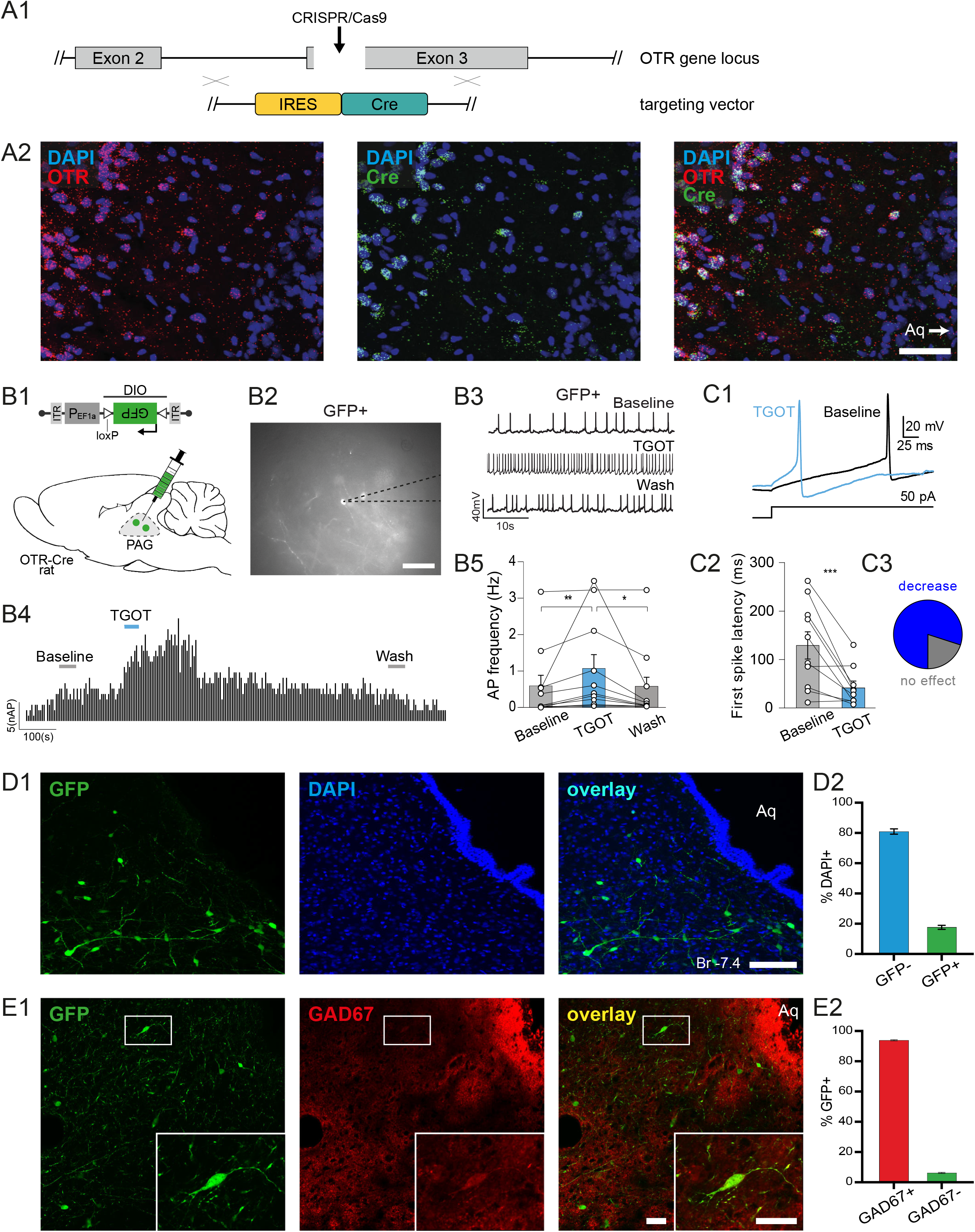
Generation of KI OTR-Cre rats and identification of vlPAG OTR neurons. **(A)** Generation and confirmation of knock-in OTR-Cre rats. **A1** Schema of the OTR gene locus and the insertion site of the targeting vector containing the IRES-Cre sequence. **A2** Images of RNA-Scope in situ hybridization showing signal from Cre (n = 47 cells) and OTR (n = 42 cells) probes. Scale bar = 200 µm **(B)** TGOT-induced increase of AP frequency of OTR neurons in the PAG. **B1** Schema of viral injection showing injection of rAAV-pEF1a-DIO-GFP in the PAG of OTR-Cre rats. **B2** Example image showing a GFP-positive cell during patch-clamp recordings. Scale bar = 20 µm. **B3** Example traces from a GFP-positive cell under baseline, TGOT application, and wash out conditions. **B4** Time course of GFP-positive cell activity (frequency distribution) upon TGOT application. **B5** Quantification of TGOT effect on AP frequency of GFP-positive cells (Baseline 0.573 ± 0.295 Hz vs TGOT 1.045 ±0.388 Hz vs Wash 0.514 ± 0.298 Hz; ** p = 0.0029, * p = 0.0168; Friedman’s test = 14.97, p < 0.0001, n = 11). Data are represented as mean ± sem and as individual paired points. **(C)** TGOT-induced change in first spike latency (FSL) of OTR neurons in the PAG. **C1**. Representative evoked currents in a GFP-positive neuron in response to a square current step (50 pA) in baseline (black line) and after TGOT application (blue line). **C2**. FSL quantification for GFP-positive neurons (OTR**^+^**): a significant difference between the baseline condition (129.31 ± 28.04 ms) and after the TGOT incubation 41.42 ± 12.37 ms, ***p = 0.0041 (paired two-tailed t test, n = 10) was found. Data are represented as mean ± sem and as individual paired points. **C3**. Proportion of neurons after TGOT incubation with a decrease of the FSL superior to 10 ms (decrease (blue) n = 8/10) or with a variation of the FSL < 10 ms (no effect (gray) n = 2/10). **(D)** Quantitative analysis of OTR cells in the PAG. **D1** Images showing GFP (green) and DAPI (blue) staining of the vlPAG from OTR-Cre rats injected with Ef1a-DIO-GFP virus. Scale bar = 100 µm. Aq = Aqueduct. **D2** Bar plot showing the percentage of vlPAG cells expressing GFP. **(E)** GAD67 staining of OTR cells in the vlPAG. **E1** Image of a vlPAG brain slice stained for GFP (green) and GAD67 (red). Scale bar = 100 µm, inset scale bar = 20 µm. **E2** Bar plot showing the percentage of GFP-positive neurons (n = 174 cells) in the vlPAG stained and not stained by GAD67 antibody.

Ex vivo electrophysiology in acute brain slices of PAG showed that application of the selective OTR agonist, [Thr^4^Gly^7^]-oxytocin (TGOT), induced a significant increase in firing of GFP+ OTR neurons which disappeared after wash out (Baseline 0.573 ± 0.295 Hz vs TGOT 1.045±0.388 Hz vs Wash 0.514 ± 0.298 Hz; n = 11; **Figure 1B1-B5**). There was no response to TGOT in recorded GFP-neurons (Baseline 0.119 ± 0.046 Hz vs TGOT 0.108 ± 0.049 Hz vs Wash 0.122 ± 0.064 Hz, n = 9, **Figure S1D1-D2**). In addition, TGOT incubation induces a significant decrease of the first spike latency (FSL) only in GFP+ OTR neurons (n = 8/10, Baseline 129.31 ± 28.04 ms vs TGOT 41.42 ± 12.37 ms, ***p = 0.0041 (paired two-tailed t test, n = 10, **Figure 1C1-C3**) and has no global effect on the FSL of GFP-OTR neurons (Baseline 31.95 ± 9.44 ms vs TGOT 37.08 ± 14.89 ms, p = 0.788 (paired two-tailed t test, n = 7, **Figure S1E1-E3**). This result shows that OTR-iRES-Cre rats correctly express functional OTRs and Cre within the same cells and that the pharmacological activation of OTRs induces a significant change in the intrinsic excitability properties of GFP+ OTR neurons.

We then quantified the number of vlPAG neurons expressing GFP and found that 396 out of 2135 (18.6%) cells were GFP-positive (**Figure 1D1-D2**), indicating that about a fifth of all vlPAG cells express OTR. Histochemical analysis of vlPAG slices further revealed that the vast majority of GFP+ cells stained positive for GAD-67, a marker of GABAergic neurons. This result indicates that virtually all of the vlPAG OTR+ neurons are GABAergic in nature (94.7%, n = 174 cells, **Figure 1E1-E2**).

### A new projection from PVN parvOT neurons to the vlPAG

To determine the origin of OT projections activating vlPAG OTR neurons, we injected a rAAV expressing mCherry under the control of the OT promotor (OTp-mCherry, (Knobloch et al., 2012) into either the PVN or SON in addition to a Cre-dependent rAAV expressing GFP (Ef1A-DIO-GFP) into the vlPAG of OTR-iRES-Cre female rats (n = 4; **Figure 2A1**). We found mCherry+ axons in close proximity to GFP+ OTR cells in the vlPAG after injection into the PVN (**Figure 2A2-A5**), but not the SON (not shown). This indicates that axons from OT neurons within the vlPAG originate exclusively from the PVN.

**Figure 2.**
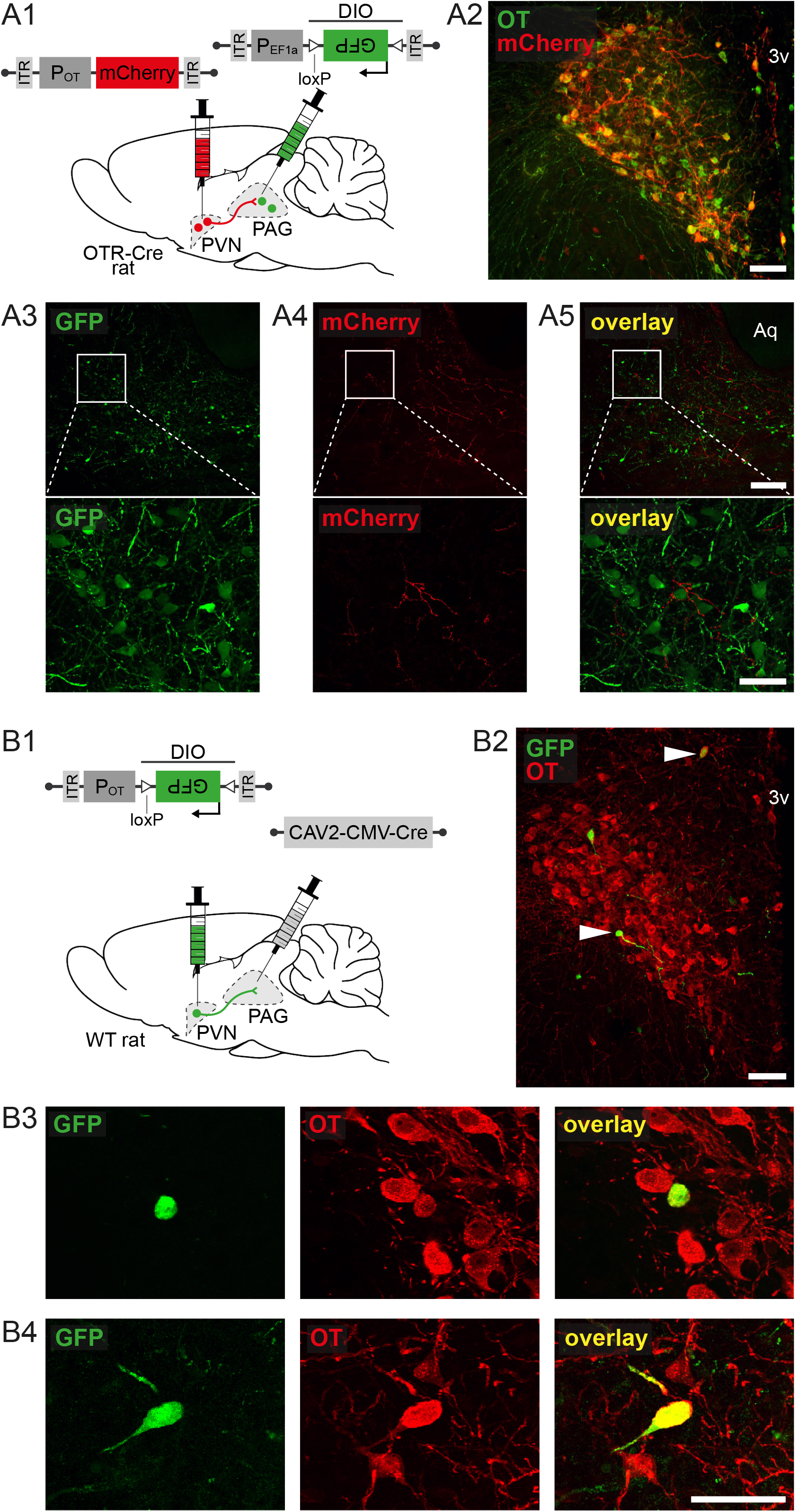
PVN ParvOT neurons send axonal projections to vlPAG. **(A)** Anterograde tracing of projections from PVN OT-neurons to the vlPAG. **A1** Schema of viral injection showing injection of rAAV-pOT-mCherry in the PVN and rAAV-pEf1a-DIO-GFP in the PAG of OTR-Cre rats (n = 4 female rats). **A2** Image showing co-localization of rAAV-pOT-mCherry and OT in the PVN. Scale bar = 200 µm. 3v = third ventricle. **A3-5** Images of GFP (green, A3) and mCherry (red, A4) staining in the vlPAG showing PVN OT fibers surrounding vlPAG GFP neurons (A5). Scale bar = 300 µm, zoom scale bar = 40 µm. Aq = Aqueduct. **(B)** Retrograde tracing of projections from PVN OT-neurons to the vlPAG. **B1** Schema of viral injection showing injection of rAAV-pOT-DIO-GFP in the PVN and rAAV-CAV2-Cre in the vlPAG of WT rats (n = 4 female rats). **B2** Image of the PVN from a rat injected with CAV2-Cre into the vlPAG and OTp-DIO-GFP into PVN, with OT stained in red. White arrows indicate co-localization of GFP and OT. Scale bar = 200 µm. **B3-4** Magnified insets of the cells indicated by white arrows in the wide field image. Scale bar = 40 µM.

We confirmed these results with retrograde tracing by injecting wild type female rats (n = 4) with CAV2-CMV-Cre into the vlPAG and rAAV-OTp-DIO-GFP into the PVN (**Figure 2B1**). We found that only a few, relatively small (n = 21 cells, 10 to 20 um diameter) OT neurons were labelled in the PVN (**Figure 2B2-B4**) whereas no labelling was detected in the SON (not shown). Furthermore, the retro-labeled cells were predominantly located at the latero-ventral edge of the PVN, although some GFP+ neurons were found in the caudal medio-dorsal region of the PVN.

Because this discrete population resembled the morphology of parvOT neurons (e.g. small size and spindle-like shape), we injected wild type female rats (n = 3) with a marker of magnOT cells, Fluorogold (FG, Santa Cruz Biotechnology, Dallas, 15 mg/kg, i.p.), to specifically label magnOT, but not parvOT neurons (Althammer and Grinevich, 2017). In parallel, the same rats received an injection of green latex Retrobeads (Lumafluor Inc., Durham, NC, USA) into the vlPAG (**Figure S2A1**) to retro-label only the OT neurons projecting to vlPAG. The histological analysis revealed that Fluorogold labelled PVN neurons did not contain the green puncta of Retrobeads in their cytoplasm (**Figure S2AB**), indicating that the OT cell projections to the vlPAG represent parvOT neurons, but not magnOT neurons.

Next, we asked whether these ParvOT → vlPAG_OTR_ neurons belonged to the same population of parvOT neurons we previously described (Eliava et al., 2016) as projecting to the SON and spinal cord (SC). We first injected wild type female rats (n = 3) with green Retrobeads into the vlPAG and red Retrobeads into the SC and found no OT+ neurons in the PVN containing beads of both colors within the same cells (**Figure S2B1-B4**). We then analyzed whether ParvOT → vlPAG_OTR_ neurons, identified in Figure 2B, are projecting axons to the SC or SON and found no detectable GFP+ axons in the cervical, thoracic or lumbar segments of the SC, nor in the SON (**Figure S2C1-C4**). Finally, we injected another cohort of wild type female rats (n = 3 per group) with CAV2-CMV-Cre into the SON (**Figure S2D1**) or the SC (F**igure S2E2**) and rAAV-OTp-DIO-GFP into the PVN. Here, we found that PVN neurons projecting to the SON also send axons to the SC, but not to the vlPAG (**Figure S2D2**) and, similarly, PVN neurons projecting to the SC also send axons to the SON, but not to the vlPAG (**Figure S2E2**). Altogether, our results indicate the existence of two distinct populations of parvOT neurons that project either to the vlPAG (present study) or to the SC (Eliava et al., 2016).

### PVN parvOT axons form direct contact with somatic and dendritic locations of vlPAG OTR neurons

Our next aim was to identify synaptic-like contacts between OT axons and OTR+ neurons in the vlPAG. First, we counted the number of OT fibers in close proximity to OTR+ neurons, relative to Bregma (**Figure 3A**-**3C**), and found a positive correlation between the two parameters (r = 0.575, p < 0.001, **Figure 3D**), indicating that OT fibers specifically target OTR neurons in the vlPAG.

**Figure 3.**
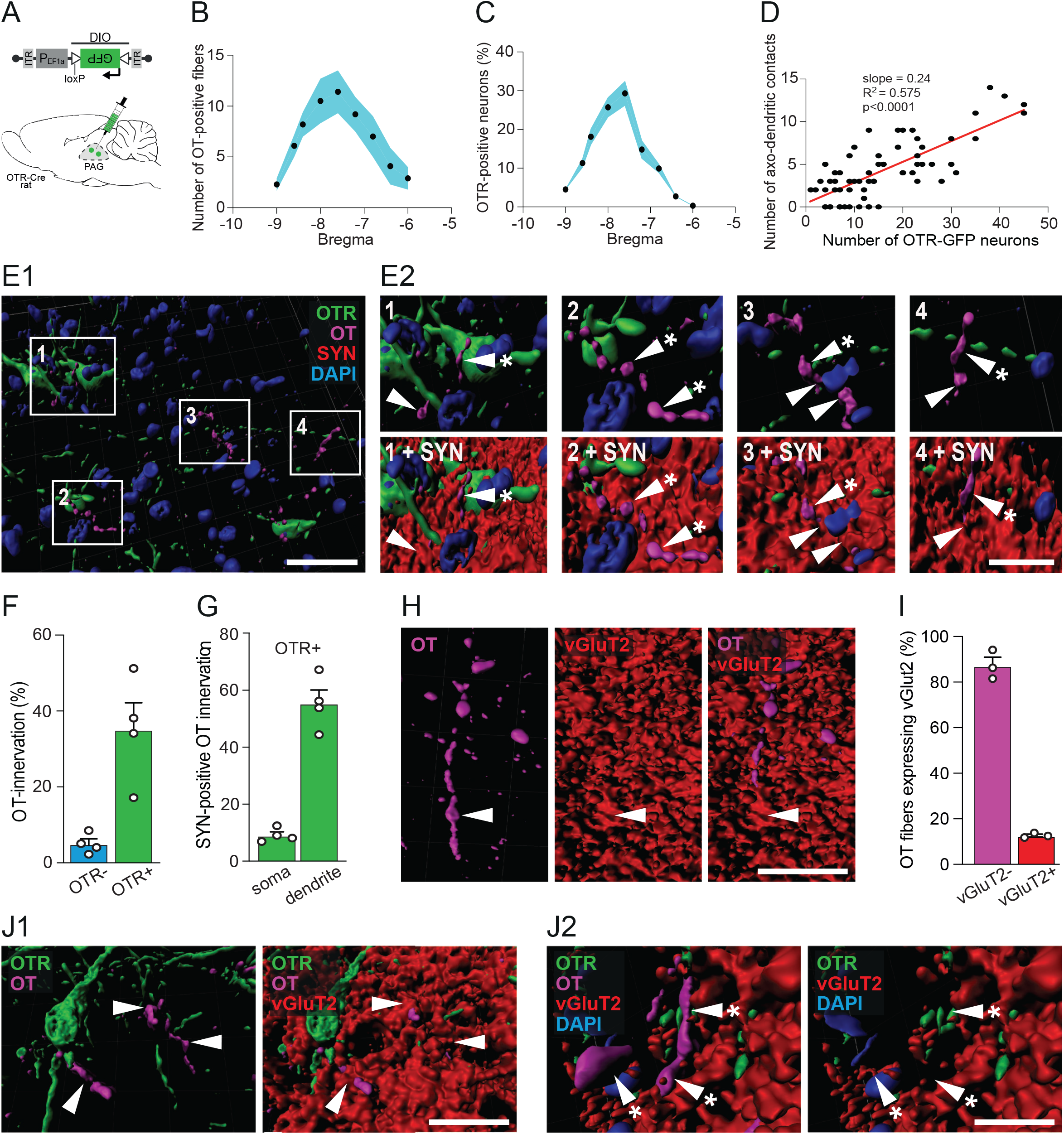
OT fibers form synaptic and non-synaptic contacts with vlPAG OTR neurons. **(A)** Schema of viral injection showing injection of rAAV-pEF1a-DIO-GFP in the PAG of OTR-Cre rats. **(B)** Graph showing the relation between the number of OT fibers in the vlPAG and the bregma level. Blue area represents SEM. **(C)** Graph showing the relation between the percentage of OTR neurons in the vlPAG and the bregma level. Blue area represents SEM. **(D)** Correlation between the number of OTR neurons and the number of OT fibers within the same slice (n = 64 slices). Each dot represents one analysed brain section. **(E)** Three-dimensional reconstruction of OTergic contacts with vlPAG OTR neurons via Imaris. **E1** Overview image showing OTR-neurons (green), OT-fibers (magenta) synaptophysin (SYN, red) and DAPI (blue). **E2** Magnified images from E1 showing contacts with or without SYN. White arrowheads indicate co-localization of OT (magenta) and SYN (red), while white arrowheads with an asterisk show a mismatch of OT and SYN. DAPI = blue, OTR = green. **(F)** Bar graph showing the percentage of OTR positive (n = 496) and negative (n = 3840) cells receiving OT innervation (< 1 µm distance between fibers and cells). n = 4, 8 images per animal, p = 0.0055. **(G)** Bar graph showings the percentage of contacts between OT and OTR-positive neurons at somatic and dendritic locations. n = 4, 8 images per animal, p < 0.0001. **(H)** Imaris reconstruction of a vGluT2-positive (red) OT fibers (magenta) within the vlPAG. **(I)** Bar graph showing that the vast majority of OT fibers within the vlPAG are vGluT2-negative (92.4%). n = 4. **(J)** 3D reconstruction of contacts between an OTR dendrite and OT fibers. **J1** Co-localization of OT (magenta) and vGluT2 (red) are indicated by white arrowheads. **J2** Mismatch of OT (magenta) and vGluT2 (red) are indicated by white arrowheads with an asterisk. DAPI = blue, OTR = green. n = 4 female rats.

Next, we injected AAV-pEf1A-DIO-GFP into the vlPAG of OTR-IRES-Cre female rats to label OTR neurons and then stained for OT, DAPI and the presynaptic marker synaptophysin (**Figure 3A**, **3E**). These sections were analyzed using Imaris software to quantify the OT innervation of GFP+ (37%) and GFP-(4%) cells (**Figure 3F**) as well as the percentage of synaptophysin-positive contacts between OT axons and GFP+ somas (7%) or dendrites (56%) (**Figure 3G**, **Figure S3A-D**). The latter indicated that the majority of OT axons predominantly form typical synaptic contacts on dendrites of PAG OTR+ neurons similar to the OT-containing synapses previously demonstrated in the brainstem and SC (Buijs, 1983).

Several reports have shown that OT is produced and released concomitantly with the conventional neurotransmitter, glutamate (Hasan et al., 2019; Knobloch et al., 2012). Therefore, we next tested whether OT-immunoreactive axons in the vlPAG also contained the glutamate transporter, vGluT2 (**Figure 3H-I**, **S3B**). This analysis revealed that only 5.3% of OT fibers were also positive for vGluT2 (**Figure 3H**). Importantly, we found synaptic-like contacts between GFP+ dendrites and both vGluT2+ (**Figure 3J1**) and vGluT2-OT fibers (**Figure 3J2**). These findings suggest that a small percentage of direct PVN OT → vlPAG contacts are glutamatergic, although the precise role of glutamate in these putative synapses remains unclear.

### Evoked OT release in the vlPAG increases neuronal activity in vivo

We then wanted to characterize the function of the PVN_OT_ → vlPAG circuit in vivo by expressing channelrhodopsin2 (ChR2) fused with mCherry (rAAV_1/2_-pOT-ChR2-mCherry; **Figure 4A**) specifically in OT neurons of the PVN (Eliava et al., 2016; Knobloch et al., 2012). In vivo PAG neuronal firing was then recorded in anesthetized female rats using silicone tetrodes coupled with a blue light (BL) that was used to stimulate PVN_OT_ axons in the vlPAG (20s at 30Hz, 10ms pulse width; **Figure 4A**, **S4**). Out of 82 recorded neurons, 21 showed an increase in firing rate (mean ± SEM; from 1.05 ± 0.39 to 17.65 ± 6.45 Hz) (**Figure 4B-E**). In contrast, two neurons, whose spontaneous activity prior to BL onset was high, showed a decreased firing rate within 300 s after the onset of BL (one cell from 25.83 to 6.95 Hz, another from 40.20 to 0.19 Hz; **Figure 4B-C**). The remaining 59 neurons did not react to BL (**Figure 4C**). We found the normalized mean activity of the excited neuronal population remained elevated for at least 300 s following the onset of BL (**Figure 4D**). Notably, the time course of spike increase was diverse, as shown by the latency (1st quantile, median, 3rd quantile) for onset (1, 4, 40.25s), peak activity (116.25, 155, 280.25s) and offset (147.75, 296, 300) (**Figure S4D**). However, the total number of active neurons was maintained throughout the 300 s period following BL stimulation (**Figure S4E**). Therefore, we conclude that BL-evoked OT release in the vlPAG leads to an overall excitation of putative OTR+ vlPAG neurons.

**Figure 4.**
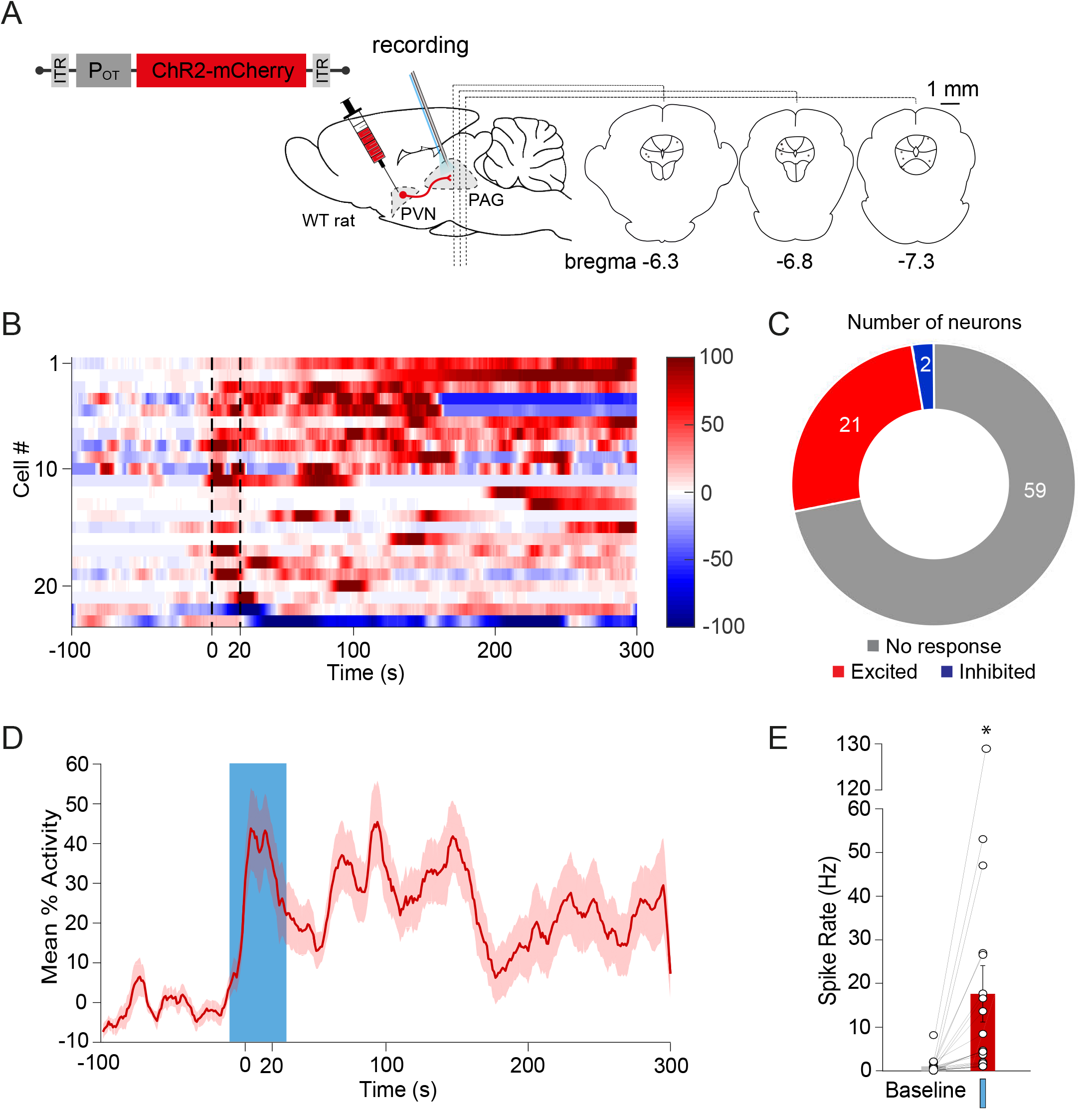
Endogenous OT release in the vlPAG increases vlPAG neuron activity. **(A)** Schema of the injection of rAAV-pOT-ChR2-mCherry in the PVN and setup for in vivo electrophysiological recordings (gray electrode), together with blue light (BL) stimulation (blue optic fiber) in the PAG. Recording site is shown on coronal drawings from anterior to posterior. **(B)** Normalized firing rate of each vlPAG neuron (n = 23) that responded to BL in the vlPAG. 473 nm of BL was added as a 10 ms pulse at 30 Hz for 20 s, 100μW/mm². Dotted lines = BL stimulation. **(C)** Recorded units’ responsiveness. Out of 82 units, the spike rates of 21 units increased (red) and 2 units decreased (blue) as a result of BL stimulation. **(D)** Mean percent activity (line) ± SEM (shaded) calculated from panel B. **(E)** Difference in mean firing rate between the period before BL (−100 to 0 s) and the maximum activity period following BL stimulation (highest value among moving means with a time window of 21 s, between 0 to +300 s after the start of BL). Paired t-test, * p < 0.05.

### Evoked OT release in vlPAG inhibits the activity of spinal cord WDR neurons in vivo

Next, we explored the downstream target of the PVN_OT_ → vlPAG_OTR_ circuit by performing in vivo BL stimulation of PVN_OT_ axons in vlPAG (vlPAG-BL) while simultaneously recording sensory wide dynamic range (WDR) neuronal responses to electrical stimulation of the hind paw receptive field (**Figure 5A**; **S5A**). We focused on WDR neurons in the spinal cord (SC_WDR_) because they are modulated by vlPAG inputs and have also been identified as an important cell population for integrating pain related signals (Ritz and Greenspan, 1985). Indeed, peripheral sensory information converges from both fast (A-beta and delta type) and slow-conducting (C-type) primary afferent fibers, which are then integrated through WDR neurons in the deep laminae of the SC. Following repetitive electric stimulation to the hind paw WDR neuron receptive field, a short-term potentiation (wind-up; WU) occurs on the synapse made by C-type fibers onto WDR neurons that causes the spike rate of the cell to reach a plateau of maximal activity (**Figure 5B1**, gray). This WU effect is typically enhanced during pain perception in animals with inflammation (Herrero et al., 2000), which suggests that WU can serve as an index of ongoing nociceptive processing (i.e. a measure of how sensitive the body is to nociceptive stimuli at a given moment). Therefore, we used WU (represented as the percentage of maximal spiking activity following electrical stimulation (1 Hz) to the hind paw receptive field) as our outcome measure for the effect of vlPAG-BL stimulation on WDR discharge, specifically from primary afferent C-fibers (**Figure S5A)**.

**Figure 5.**
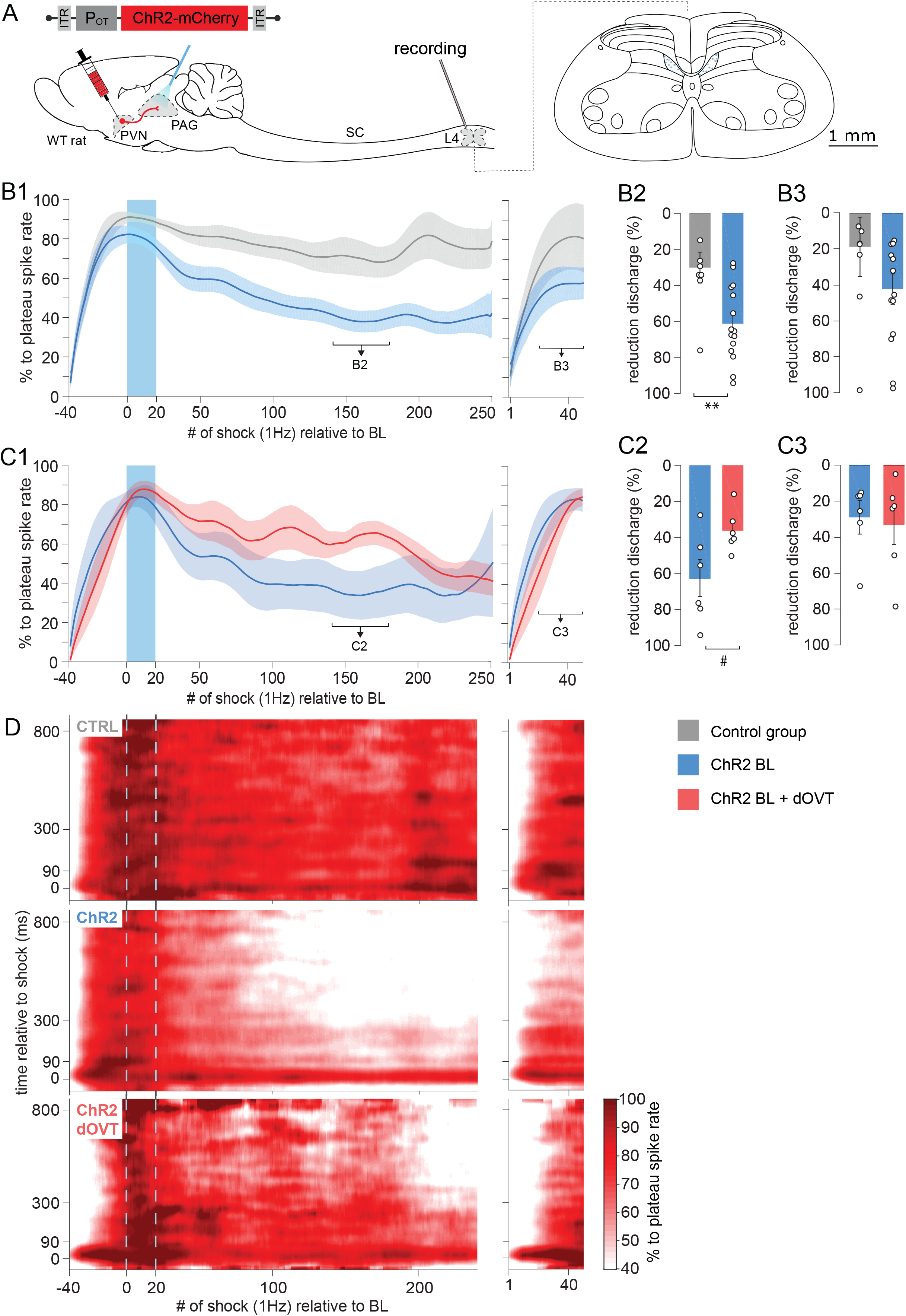
Endogenous OT release in vlPAG reduces WDR spinal cord neuronal activity. **(A)** Schema of the injection of rAAV-pOT-ChR2-mCherry (or rAAV-pOT-mCherry for control group) in the PVN and setup for in vivo electrophysiological recordings (gray electrode) of WDR neurons in the rat spinal cord (SC) at the lumbar 4 (L4) level during optogenetic BL stimulation (blue optic fiber) in the vlPAG to activate ChR2-expressing axons originating from PVN OT neurons. Recording sites in layer 5 are shown in the coronal drawing of L4. **(B)** vlPAG BL effect on the spike rate of WDR’s C-fiber discharge. (**B1**) Mean time course observed after vlPAG BL in control rats (gray, n = 8) and OT ChR2-expressing rats (blue, n = 14). Left and right panels show two consecutive recordings separated by 300 s. Line shadows represent SEM. (**B2**) Percentage of reduction expressed as the minimum level activity observed after a wind-up plateau phase, 140 – 180 s and (**B3**) 570 – 600 s after BL onset. Unpaired Wilcoxon rank sum test; ** p < 0.01. **(C)** PAG OTR contribution to the vlPAG BL effect on the spike rate of WDR’s C-fiber discharge. (**C1**) Mean time course observed after vlPAG BL in OT ChR2-expressing rats (blue, n = 6), and in the same recordings after dOVT injection in the vlPAG (red, n = 6). Left and right panels show two consecutive recordings separated by 300 s. Line shadows represent SEM. (**C2**) Percentage of reduction expressed as the minimum level activity observed after a wind-up plateau phase, 140 – 180 s and (**B3**) 570 – 600 s after BL onset. Paired Wilcoxon signed rank test; # p < 0.01. **(D)** Mean smoothed raster plot of WDR discharge level along the relative timing to each single electric shock on the hind paw (vertical axis) and along the accumulating trials of electric shock (horizontal axis), in CTRL animals (top, n = 8), ChR2 animals (middle, n = 14), and ChR2 animals after dOVT injection in PAG (bottom, n = 6).

The results show that, prior to any vlPAG-BL stimulation, all cells exhibited the maximal WU effect 30 s after the onset of electrical stimulation (**Figure 5B1**). In control animals (CTRL) that received vlPAG-BL in the absence of ChR2 expression, the WU remained stable up to 250 s after the plateau, despite a gradual reduction over time that was not statistically significant. In contrast, vlPAG-BL stimulation in animals expressing ChR2 in OT neurons showed a significant decrease in WU compared to control animals (**Figure 5B1, B2**; CTRL 30.12 ± 8.60 vs ChR2 61.28 ± 5.37 %, p = 0.0074, n = 8 and 14, respectively). This trend was maintained up to 600 s after the end of vlPAG-BL stimulation, as seen in a second series of recordings of the same neurons (**Figure 5B1**, right panel, **Figure 5B3**). While the magnitude of WU reduction was significantly larger in ChR2 animals than CTRL animals, there was no difference in the “inflection” timing of WU dynamics. Specifically, there was no significant difference between CTRL and ChR2 animals for latency (s) to reach the maximum WU (**Figure S5B**). Importantly, all recorded WDR neurons were impacted by vlPAG-BL, highlighting the effectiveness of this circuit in gating the nociceptive signal at the spinal cord level.

In order to confirm that the recorded effect on WU was due to OT release in the vlPAG, we ran a second series of experiments in which we allowed the WU effect to dissipate over a 10 m interval following the initial stimulation protocols in the ChR2 group. We then infused the specific OTR antagonist, [d(CH2)5,Tyr(Me)2,Orn8]-vasotocin (dOVT), into the vlPAG prior to repeating the same protocol described above. We found that dOVT infusion caused a significant reduction of the vlPAG-BL stimulation effect such that WU was greater when dOVT was onboard than it was during the initial stimulation protocol (ChR2 + dOVT: 36.27 ± 4.80 %, p = 0.0313, n = 6) (**Figure 5C1**, **C2**). **Figure 5D** summarizes the vlPAG-BL effect in the different groups during the period of maximum WU reduction (from 140 to 180 s), since dOVT lost its effectiveness after this period (**Figure 5C1**, **C3**), possibly due to diffusion out of the PAG region.

### OT in the vlPAG induces analgesia during both inflammatory and neuropathic pain

Because the vlPAG is known to be a key component of an important descending pain modulatory system, and OT is known to exert an analgesic effect (Yang et al., 2011), we hypothesized that OTR+ neurons in the vlPAG are involved in pain processing. To test this hypothesis, we injected rAAV-pEF1A-DIO-GFP into the vlPAG of male OTR-iRES-Cre rats. The rats were subdivided into four groups (n = 7-8 per group) :1) no manipulation (Control), 2) inflammatory-induced pain sensitization after complete Freund adjuvant injection in the posterior right paw (CFA), 3) acute mechanical nociception (Pain), 4) mechanical nociception 24h following hind paw CFA injection (CFA + Pain). Rats were euthanized 30 m following each manipulation and brain sections containing vlPAG were collected. We next used c-Fos staining to compare the number of recently activated OTR (GFP+) expressing cells across the groups (**Figure 6A-B**). We found that rats exposed to either painful stimuli or painful stimuli following CFA injection had significantly more activated OTR neurons in the vlPAG (38.1 ± 9.2% and 35.9 ± 3.4%, respectively; p < 0.01) than rats not exposed to nociceptive stimulation (18.4 ± 9.6%). There was no statistically significant difference between the CFA group (26.0 ± 6.8%) and the control group (**Figure 6A**).

**Figure 6.**
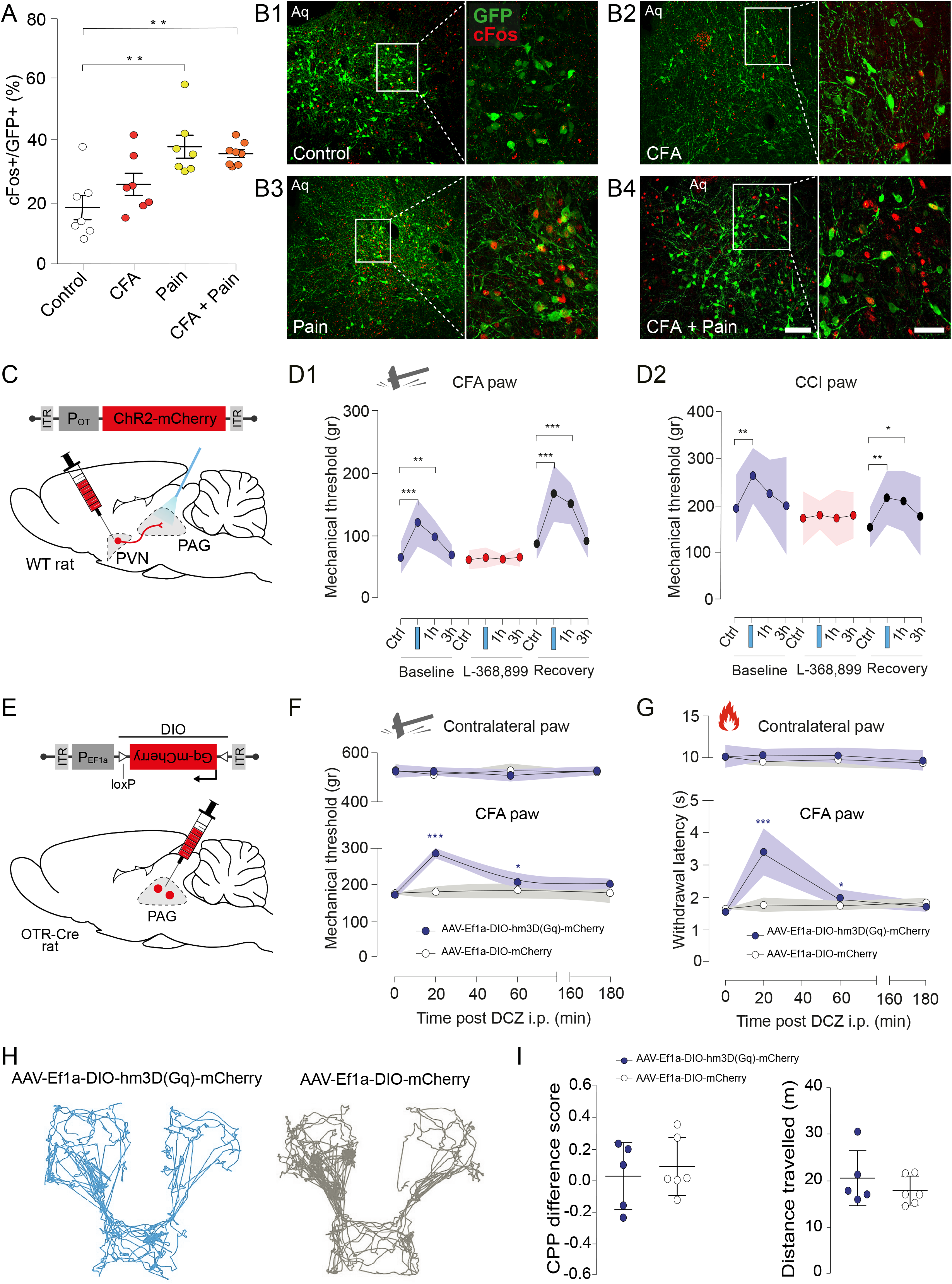
Evoked OT release in vlPAG reduces mechanical hyperalgesia. **(A)** Percentage of c-Fos positive vlPAG OTR neurons under the control condition, painful stimulation, CFA inflammation, and painful stimulation combined with CFA. n = 7-8 per group, Kruskall Wallis test H = 12.01, p < 0.01, CTRL vs pain and CTRL vs pain + CFA p < 0.05, CTRL vs CFA p > 0.05. All results are expressed as average ± SEM. **(B)** Examples of images showing c-Fos (red) and GFP (green) staining of vlPAG under the different experimental conditions (**B1-4**). Scale bar = 200 µm, inset scale bar = 75 µm. Aq = aqueduct. **(C)** Schema of the injection of rAAV-pOT-ChR2-mCherry in the PVN and optic fiber implantation in the PAG. **(D)** Threshold of mechanical pain was raised by PAG-BL. The effect of vlPAG-BL was measured at 5 min, 1 h and 3 h after vlPAG-BL for the CFA-injected paw (**D1**) as well as the contralateral paw (**D2**). Blue background represents BL stimulation. **** p < 0.0001, n = 10. **(E)** Schema of the injection of rAAV-pEf1a-DIO-Gq-mCherry in the PAG. **(F)** Mechanical pain threshold after DCZ administration in the CFA and contralateral paw of female rats expressing Gq-mCherry (blue) or mCherry only (gray) in vlPAG OTR neurons. All results are expressed as average ± SEM. * p < 0.01, *** p < 0.0001, n = 5-6 per group. **(G)** Thermal pain threshold of female rats expressing Gq-mCherry (red) or mCherry only (gray) in vlPAG OTR neurons after DCZ administration during normal or inflammation (CFA) conditions. All results are expressed as average ± SEM. * p < 0.01, *** p < 0.0001, n = 5-6 per group. **(H)** Representative activity traces during the CPP test. **(I)** Graphs showing the ΔCPP score (left) and total distance travelled (right) for the test and control groups. n = 6 per group

To verify the functional significance of vlPAG projecting OT neurons in the processing of inflammatory pain, a separate cohort of female wild type rats received vlPAG injections of rAAV_1/2_-pOT-ChR2-mCherry. We then compared the within-subject effect of optogenetically-evoked OT release in the vlPAG (vlPAG-BL) on mechanical pain sensitivity both with and without the presence of CFA-induced inflammatory hyperalgesia (**Figure 6C-D**). We found that vlPAG-BL stimulation significantly, but not entirely, alleviated CFA-induced hyperalgesia as indicated by an increase in the mechanical pain threshold from 68.77 ± 9.39 g to 124.89 ± 12.76 g (**Figure 6D1**; n = 10). Injection of the blood-brain barrier (BBB)-permeable OTR antagonist, L-368,899, completely blocked the effect of BL in the vlPAG (from 62.75 ± 5.31 g to 71.99 ± 5.14 g, **Figure 6D1**; n = 10). After complete wash out of L-368,899, the effect of vlPAG-BL returned to its baseline level (from 92.95 ± 8.89 g to 179.40 ± 19.13 g; **Figure 6D1**; n = 10). Finally, we found that vlPAG-BL had no effect on mechanical sensitivity in the absence of any peripheral sensitization when testing the contralateral paw (**Figure 6D2**).

To confirm that this effect was driven by vlPAG OTR neurons, we injected male and female OTR-Cre rats with viruses containing either an excitatory chemogenetic receptor, rAAV-Ef1a-DIO-hm3D(Gq)-mCherry, or a control virus rAAV-Ef1a-DIO-mCherry (n = 5-6 per group) (**Figure 6E**, **S6B**). We then repeated the CFA-induced inflammatory hyperalgesia experiments described above and found that chemogenetic excitation of vlPAG OTR neurons by i.p deschloroclozapine (DCZ) induced a significant increase in mechanical pain threshold from 171.3 ± 12.5 g to 285.5 ± 14.2 g (**Figure 6F**). This effect was not observed in the contralateral paw of the same animals nor in the control virus group that received DCZ injection (**Figure 6F**). A similar effect was found in the hot plate test of thermal pain sensitivity, where DCZ increased the latency of response to thermal stimuli from 1.56 ± 0.13 s to 3.41 ± 0.73 s (**Figure 6G**), this was again not found in the contralateral paw of the same animals nor in the control group (**Figure 6G**). Finally, we repeated these experiments in male rats and found the same results (**Figure S6C**, **S6D**, n = 6 per group), ruling out a potential sexual dimorphism of this circuit.

We next sought to test if this novel PVN_OT_ → vlPAG_OTR_ circuit is involved in neuropathic pain, given that other pain-related OT pathways fail to affect such symptoms (Eliava et al., 2016; Wahis et al., 2021). To address this, we performed vlPAG-BL stimulation in the chronic constriction of the sciatic nerve (CCI) model of neuropathic pain (Austin et al., 2012). We found that BL-evoked OT release in the vlPAG increased the threshold of response to mechanical stimuli from 195.7 ± 72.3 g to 265.3 ± 58.9 g (**Figure S6A1-2**; n = 7). Similar to the previous experiment, injection of L-368,899 completely blocked the effect of BL in the vlPAG (from 174.6 ± 56.4 g to 181.2 ± 29.7 g, **Figure S6A2**; n = 7) and the effect of vlPAG-BL was fully restored after wash out of L-368,899 (from 155.5 ± 43.1 g (baseline) to 218.3 ± 57.5 g (BL); **Figure S6A2**; n = 7). In addition, vlPAG-BL did not modulate the mechanical pain threshold of the contralateral paw (**Figures S6A3**).

Finally, to understand whether the analgesia was caused by a modulation of the sensory and/or emotional component of pain (King et al., 2009), we performed a conditioned place preference test using the same cohort from the chemogenetic experiments above (**Figure S6E**). The animals’ baseline chamber preference was determined during habituation (see Methods for details) and used as the saline-paired control chamber. In contrast, the innately non-preferred chamber was paired with DCZ in order to stimulate vlPAG OTR cells expressing hm3D(Gq) (**Figure 6E**). Analysis of the ratś behavior on test day revealed no significant change in preference for the DCZ-paired chamber (**Figure 6H-I**). Importantly, this was not due to an effect on locomotion as there were no differences between the test and the control group in the total distanced travel during the experiment (**Figure 6I**).

## Discussion

Here we describe a new analgesic pathway recruited by newly discovered parvOT neurons projecting to the vlPAG (**Figure 1-2**), where they activate GABAergic OTR expressing neurons (**Figure 3-4**), which then leads to a decreased response to nociceptive stimuli in spinal cord WDR neurons (**Figure 5**). We further showed that activation of this novel circuit specifically reduces pain sensation (**Figure 6**), without alteration of the affective component of pain.

A vast amount of literature has shown that OT exerts analgesic effects by acting on various targets of pain-associated areas in the central and peripheral nervous systems (Boll et al., 2018; Poisbeau et al., 2017). The contribution of OT to analgesia is generally attributed to two pathways. The first is an ascending OT pathway that modulates the activity of brain regions processing the emotional and cognitive components of pain, such as the amygdala, in which OT alleviates anxiety, especially in the context of chronic pain in rodents (Hasan et al., 2019; Knobloch et al., 2012; Wahis et al., 2021). In humans, OT was found to decrease neural activity in the anterior insula with repeated thermal pain stimulation, thereby facilitating habituation to the cognitive component of the painful stimuli (Herpertz et al., 2019). The second is a descending OT pathway that indirectly promotes analgesia by reducing the activity of SC WDR neurons, which relay nociception in response to painful stimuli (DeLaTorre et al., 2009; Eliava et al., 2016). While this specific descending OT pathway is effective, it is restricted to inflammatory pain (Eliava et al 2016). In contrast, here we described a powerful descending OT pathway that is effective for both inflammatory and neuropathic pain across thermal and mechanical modalities.

### A novel and distinct parvOT neuronal population promotes analgesia via the vlPAG

We identified a novel population of parvOT neurons that projects to the vlPAG, but not the SON nor the SC. Furthermore, we found that these neurons form synapses with little contribution of glutamate, thus supporting the idea of local axonal delivery of OT, as opposed to volume transmission (Chini et al., 2017; Landgraf and Neumann, 2004; van den Pol, 2012).

Consistent with previous reports showing that electrical stimulation of the PAG inhibits the firing of dorsal horn neurons in the SC (Liebeskind et al., 1973) and generates analgesia (Basbaum et al., 1977), we found that nociceptive transmission from C-type primary afferents to WDR neurons in the SC was effectively repressed by endogenous OT release in the vlPAG. Notably, this effect peaked 250 s after OT release and was still observed 10 minutes after the cessation of blue light. This finding suggests that OT triggers a lasting activation of OTR expressing cells in the vlPAG, which then continually drives the regulation of SC WDR neurons for several minutes after initial OT release. Indeed, in a separate set of experiments, we showed that exogenously applied OTR agonist as well as endogenously evoked OT release excites neurons of the vlPAG and induces analgesia in the context of inflammatory pain. Notably, optogenetic release of OT in the vlPAG led to activation of individual neurons at various times, mostly within the first 40 seconds after the onset of blue light stimulation. The neurons’ offset timings were also diverse and usually lasted for several minutes after the offset of blue light stimulation. The reason for such variability in the offset times is still unclear. One possibility may stem from the G-protein coupled metabotropic receptor nature of OTRs, which typically produce “slow” post synaptic currents lasting on the order of minutes (Knobloch et al., 2014). Furthermore, although OT axons in the midbrain make synapses (Buijs, 1978, 1983), direct release of OT into the synaptic cleft has never been demonstrated. Thus, it is more likely that OT diffuses from axonal terminals or axonal varicosities *en passant* in the vicinity of OTR neurons (Chini et al., 2017). In which case, the action of OT could be synergized across multiple OTR expressing cells, resulting in long-lasting excitation driven by the sum of different active timings. This idea leads to another possible mechanism in which the OT-induced modulation of this pathway relies on the influence of additional non-neuronal cell types within the network, such as astrocytes, as was previously shown in the amygdala (Wahis et al., 2021). While the specific mechanisms behind the lasting analgesic effect of OT release in the vlPAG are unclear, they would certainly play a critical role in its development as a potential therapeutic target and, therefore, warrant future research.

Importantly, we confirmed the uniqueness of this parvOT pathway by showing that the previously identified parvOT → SC_WDR_ and parvOT → SON pathways do not project to vlPAG. Interestingly, the level of reduction in nociception caused by optogenetic stimulation of parvOT neurons projecting to SON (Eliava et al., 2016) resembled the effect of stimulating vlPAG OT axons. Redundant, parallel circuits that play identical roles in the brain have been previously described (e.g. for feeding behavior (Betley et al., 2013)). Therefore, the direct projections from parvOT neurons to the SC and the indirect influence of parvOT neurons on sensory WDR neurons via the vlPAG can be interpreted as parallel circuits capable of independently promoting analgesia, particularly in the context of inflammatory pain. Although activation of both circuits results in similar electrophysiological inhibition of WDR neurons, they could be triggered by different situations, at different time points, or in different painful contexts. Indeed, we found that the PVN_OT_ → vlPAG_OTR_ circuit promotes analgesia in the chronic neuropathic pain condition and in both male and female rats, whereas previous work found that the parvOT → SC_WDR_ circuit does not (Eliava et al., 2016). Moreover, considering that the vlPAG is an important area for the regulation of various defensive behaviors (Tovote et al., 2015), and that OT can mediate defensive behaviors, it will be important for future research to determine if vlPAG OTR neurons might be involved in other functions beyond nociception.

### vlPAG OTR neurons reduce pain sensation, but not its affective component

We further found that painful stimulation increased c-fos levels in vlPAG OTR neurons, indicating that these neurons are endogenously recruited in the context of pain processing. These findings confirm and expand on previous work demonstrating that PAG OTR neurons in mice express c-fos after noxious stimulation (Saito et al., 2021).

By using our newly generated OTR-IRES-Cre transgenic rats, we were able to specifically activate OTR neurons in the vlPAG. We found that this activation led to a decrease in nociception in both inflammatory and neuropathic pain, but failed to alter place preference. This suggests the circuit we dissected here is not involved in the affective, memory component of pain (King et al., 2009). Interestingly, the opposite effect of OT on emotional valence, but not on physical pain perception has been demonstrated in the central nucleus of amygdala (Wahis et al., 2021). In addition, OT was reported to modulate pain at the level of the insular cortex by enhancing GABAergic transmission and causing downstream effects leading to reduced nociceptive signaling in the spinal cord (Gamal-Eltrabily et al., 2020). Taken together, these findings further emphasis the sheer variety of effects mediated by OT and highlight the need for continued efforts to dissect the precise anatomical and functional characteristics of the central OT system.

In conclusion, we identified a new subpopulation of parvocellular OT neurons that mediate analgesia by recruiting the PAG-controlled descending pain modulatory system (**Figure 7**). This study further describes and supports the role of OT as an analgesic molecule and points to the OTR as a potential therapeutic target. To this end, we generated the OTR-iRES-Cre line of rat, which will greatly enhance our ability to research this therapeutically relevant receptor. Finally, it should be noted that the inconsistent results regarding sex differences and subjective pain ratings (Pfeifer et al., 2020) found in human clinical studies of OTs effects on pain (Boll et al., 2018) may be due to the limitations of intranasal OT administration (Leng and Ludwig, 2015). Therefore, future research should be oriented towards developing synthetic OT agonists with the ability to cross the blood brain barrier more efficiently than OT itself (Busnelli and Chini, 2017; Busnelli et al., 2016; Hilfiger et al., 2020; Muttenthaler et al., 2017).

**Figure 7.**
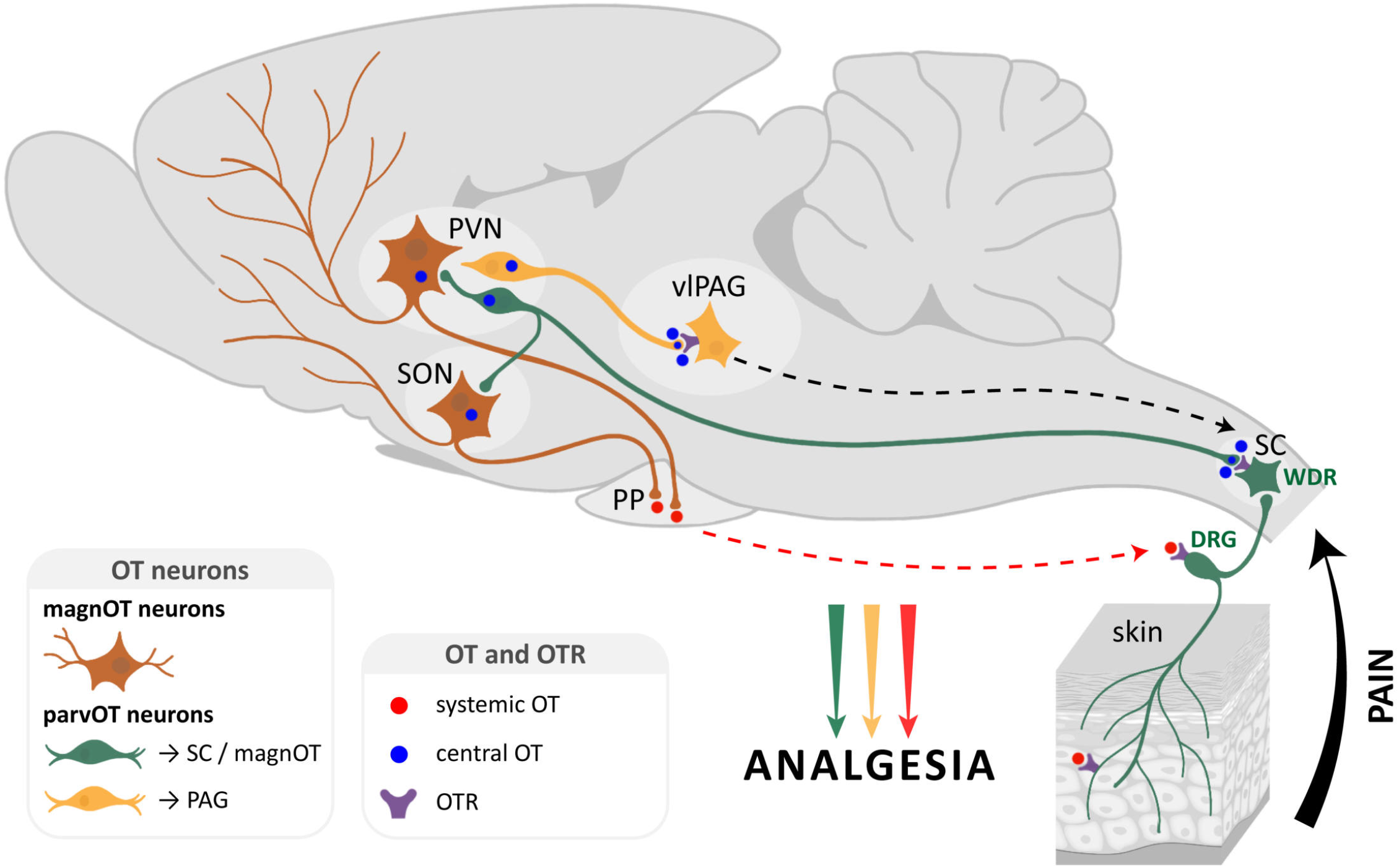
Two distinct ParvOT neuronal populations promote analgesia via release of OT in the vlPAG and in the blood and spinal cord. We hypothesize that two parallel parvOT pathways are activated by pathological, painful stimuli. Both pathways release OT in various brain regions and the periphery which then leads to a reduction in nociception.

## Acknowledgments

This work was supported by the University of Strasbourg Institute for Advanced Study (USIAS) fellowship 2014-15, Fyssen Foundation research grant 2015, NARSAD Young Investigator Grant 24821, Agence Nationale de la Recherche (ANR, French Research Foundation) grants n° 19-CE16-0011-0 and n° 20-CE18-0031 (to AC); ANR-DFG grant GR 3619/701, PHC PROCOPE and PICS07882 grants (to AC and VG); Deutsche Forschungsgemeinschaft (DFG, German Research Foundation) grants GR 3619/15-1, GR 3619/16-1(to VG); SFB Consortium 1158-2 (to VG, SH and BD); French Japanese governments fellowship B-16012 JM/NH (to MI); Fyssen Foundation fellowship (to AL); Région Grand Est fellowship (to DK); DFG Postdoc Fellowship AL 2466/1-1 (to FA); DAAD Postdoc Short term research grant 57552337 (to RP); DFG Walter Benjamin Position – Projektnummer 459051339 (to QK). National Heart, Lung, and Blood Institute Grant NIH HL090948, National Institute of Neurological Disorders and Stroke Grant NIH NS094640, and funding provided by the Center for Neuroinflammation and Cardiometabolic Diseases (CNCD) at Georgia State University (to JES). The authors thank Romain Goutagny and Vincent Douchamps for in vivo electrophysiology advices, the Chronobiotron UMS 3415 for all animal care and the technical plateau ComptOpt UPR 3212 for behavior technical assistance.

## Authors contributions

Conceptualization, AC; Methodology, AC, AL, DK, FA, LH, MI, MM, OL, QK, VG; Analysis, AL, DK, FA, LH, MI; in situ hybridization, FA; Immunohistochemistry, AL, FA, SK; Ex vivo patch-clamp electrophysiology, DK, LH; In vivo electrophysiology, MI, MM, OL; Behavior LH, MI; transgenic rats line generation, KS, DB; Writing, AC, AL, BD, FA, JS, MI, OL, PD, RP, SH, SK, VG; Funding acquisition AC, VG; Supervision AC, VG; Project administration, AC.

## Materials and Methods

### Animals

Adult female and male Wistar wild type and Sprague Dawley OTR-IRES-Cre rats (>8 weeks old; 250 - 350 g; Chronobiotron, Strasbourg, France) were used for this study. All animals were tattooed, sexed and genotyped (Kapa2G Robust HotStart PCR Kit, Kapa Biosystems; Hoffman La Roche) one week after birth. Animals were housed by sex, in groups of three under standard conditions (room temperature, 22 °C; 12 / 12 h light / dark cycle) with ad libitum access to food, water and behavioral enrichment. All animals that underwent behavioral testing were handled and habituated to the experimenter two weeks before stereotaxic surgery. After one week of post-surgical recovery, the rats were habituated to the applicable behavioral testing room and handling routines for an additional two weeks prior to the start of experiments. All behavioral tests were conducted during the light period (i.e., between 7:00 and 19:00). All experiments were conducted in accordance with European law, under French Ministery license 3668-2016011815445431 and 15541-2018061412017327, and German Animal Ethics Committee of the Baden Württemberg license G-102/17.

### Viral cloning and packaging

Recombinant Adeno-associated virus (serotype 1/2) carrying either a conserved region of the OT promoter or Ef1A promoter and genes of interest in direct or “DIO” orientations were cloned and produced as reported previously (Knobloch et al., 2012). Briefly, HEK293T cells (#240073, Addgene, USA) were used for viral production. rAAVs produced included: rAAV-OTp-mCherry(or Venus), rAAV-OTp-ChR2-mCherry, rAAV-Ef1A-DIO-GFP (or mCherry), and rAAV-Ef1A-DIO-hM3Dq-mCherry. The canine adenovirus serotype 2 (CAV2-CMV-Cre) was purchased from the Institute of Molecular Genetics in Montpellier CNRS, France. rAAV genomic titers were determined with QuickTiter AAV Quantitation Kit (Cell Biolabs, Inc., San Diego, California, USA) and RT-PCR using the ABI 7700 cycler (Applied Biosystems, California, USA). rAAVs titers were between 10^9^ - 10^10^ genomic copies/μl.

### Stereotaxic injections

All surgeries were performed on rats anesthetized with 2.5% isoflurane and receiving Bupivacaine (s.c., 2mg/kg) or carprofen (i.p., 5mg/kg) and lidocaine applied locally (Tang et al., 2022). rAAVs were injected into the PVN, SON and vlPAG in different combinations, as needed by each experiment, and allowed to express for four weeks. The coordinates were chosen using the Paxinos rat brain atlas (Paxinos et al., 1985)(PVN: ML: +/-0.3 mm; AP: -1.4 mm; DV: -8.0 mm; SON: ML: +/-1.8 mm; AP: -1.2 mm; DV: -9.25 mm; PAG: ML: +/-0.5 mm; AP: -7.0 mm; DV: -5.9/-5.0 mm). Each site was injected with a total of 300 nL of viral solution via a glass pipette at a rate of 150 nl/min using a syringe pump. Verification of injection and implantation sites, as well as expression of genes of interest were confirmed in all rats post-hoc (see “Histology” section). Rhodamine conjugated Retrobeads (Lumafluor Inc., Durham, NC, USA) were diluted 1:10 with 1x PBS and injected at a volume of 150 nl. Spinal cord Retrobeads injections was performed during the same surgery as virus injection (See “In vivo extracellular recording of WDR SC neurons” for details on the spinal cord surgery).

### Generation of OTR-IRES-Cre rats

#### Cloning of the rat OXTR-Cre targeting vector

The OXTR-Cre targeting vector was cloned by modifying the plasmid Snap25-IRES2-Cre (Allen Institute for Brain Science (Harris et al., 2014)) The final vector contained the iRES2-Cre sequences, followed by a bovine growth hormone polyadenylation site, and homology arms for targeted integration of the oxytocin receptor locus, comprised of 1.3 kb and 1.4 kb genomic sequences. Homology arms were generated by PCR on genomic DNA from Sprague Dawley rats using the following primer pairs: OXTR_fwd_upper (5’ GTCGACAGAAAACTGGTGGGTTTGCC 3’) together with OXTR_rev_upper (5’ GCTGCTAGCGAAGACTGGAGTCCACACCACC 3’) and OXTR_fwd_lower (5’ ACCCGGGAATTCTGTGCATGAAGCTGCATTAGG 3’) together with OXTR_rev_lower (5’ TAGTTTAAACGTGCATTCGTGTATGTTGTCTATCC 3’). The upper homology arm was inserted using the restriction enzymes SalI and NheI, while the lower arm was inserted using XmaI and PmeI. Vector sequences can be obtained upon request.

#### Design of gRNAs and functional testing

We used the online tool CRISPOR (http://crispor.org) for selection of the guide RNA (gRNA) target sites in the OXTR gene (Haeussler et al., 2016). Three gRNA target sites were chosen with high specificity scores (> 83, (Hsu et al., 2013)) for binding in the OXTR 3’ UTR site where we aimed to introduce the IRES-Cre coding sequences.

For identification of the most effective gRNA, dual expression vectors based on px330 were cloned, harboring an expression cassette for the selected gRNAs and Cas9. In addition, the OXTR 3’ UTR gRNA target regions were inserted into a nuclease reporter plasmid (pTAL-Rep, (Wefers et al., 2013)) in between a partly duplicated, nonfunctional β-galactosidase gene. Hela cells were transfected with a combination of one of the px330 plasmids and the OXTR specific reporter vector. After transfection, the Cas9-nuclease-induced double-strand breaks stimulated the repair of the lacZ gene segments into a functional reporter gene, the activity of which was determined in cell lysates using an o-nitrophenyl-β-D-galactopyranosid (ONPG) assay. A luciferase expression vector was also added to the transfection mix and luciferase activity was measured for normalization. The most effective gRNA target site including PAM was determined as 5’ ACTCCAGTCTTCCCCCGTGG**TGG** 3’

#### Specificity of the CRISPR/Cas induced genomic modification

The CRISPOR program was used to identify potential off-target sites for the selected OXTR gRNA (5’ACTCCAGTCTTCCCCCGTGGTGG 3’). No off-target sites were detected in an exonic sequence or on the same chromosome. In addition, only two potential target sites harboring at least four mismatches in the 12 bp adjacent to the PAM could be identified by the software.

#### Generation of transgenic rats

Sprague Dawley (SD) rats (Charles River) were bred in standard cages (Tecniplast) under a 12-h light/dark cycle in a temperature-controlled environment with free access to food and water at the Central Institute of Mental Health, Mannheim. All animal protocols were approved by the Regierungspräsidium Karlsruhe. SD single-cell embryos were injected using standard microinjection procedure (Shao et al., 2014). In brief, microinjections were performed in the cytoplasm and male pronuclei of zygotes with a mixture of Cas9 mRNA (10 ng·/μl), sgRNA expression vector (6 ng/μL) and the OXTR-Cre targeting vector (2 ng/μl) as the repair substrate. The injected embryos were cultured in M2 Medium at 37°C in 5% CO2 and 95% humidified air until the time of injection. Surviving oocytes were transferred to the oviducts of pseudo pregnant Sprague Dawley rats. Transgenic animals were identified by polymerase chain reaction of tail DNA (DNeasy kit, Qiagen, Hilden, Germany) using primers for Cre (Schönig et al., 2002).

DNA of Cre positive animals was further used for detection of homologous recombination at the OXTR locus. For this purpose, the targeted region was amplified by PCR using the Q5 polymerase (NEB) with 100 -200 ng of genomic DNA as a template. Primers were selected which bind both up and downstream of the insertion site (outside of the homology arms), and were each combined with a primer located within the IRES2-Cre construct. For the 5’ insertion site, the primers OXTR_check (5′ CAGCAAGAAGAGCAACTCATCC 3′) together with Cre_rev (5′ CATCACTCGTTGCATCGAC 3′) and, for the 3’ site, CRISPR_bGH_fwd (5’ GACAATAGCAGGCATGCTGG 3’) together with OXTR_rev_check (5’ AGCCAGGTGTCCAAGAGTCC 3’) were used.

### Histology

After transcardial perfusion with 4% paraformaldehyde (PFA) and post fixation overnight, brain sections (50 µm) were collected by vibratome slicing and immunohistochemistry was performed as previously described (Knobloch et al., 2012). The list of primary antibodies used is available in the key resource table. For secondary antibodies, signal was enhanced by Alexa488-conjugated IgGs (1:1000) or CY3-conjugated or CY5-conjugated antibodies (1:500; Jackson Immuno-Research Laboratories). All images were acquired on a confocal Leica TCS microscope. Digitized images were processed with Fiji and analyzed using Adobe Photoshop. For the visualization of OTergic axonal projections within the PAG, we analyzed brain sections ranging from bregma -6.0 to -8.4 mm.

### RNAScope in situ hybridization

RNAScope reagents (Advanced Cell Diagnostics, Inc., PN320881) and probes (OTR: 483671-C2 RNAscope Probe and Cre: 312281 RNAscope Probe) were used to detect the presence of specific mRNA expression using in situ hybridization. Brains were processed as described above using nuclease-free PBS, water, PBS and sucrose. We followed the manufacturer’s protocol with a few modifications: 1) immediately after cryosectioning, slices were washed in nuclease-free PBS to remove residual sucrose and OCT compound. 2) Hydrogen peroxide treatment was performed with free-floating sections prior to slice mounting. 3) Sections were mounted in nuclease-free PBS at room temperature. 4) Pretreatment with Protease III was performed for 20 minutes at room temperature. 5) No target retrieval step was performed.

### Three-dimensional reconstruction and analysis of OT-OTR contacts in the PAG

Confocal images were obtained using a Zeiss LSM 780 confocal microscope (1024x1024 pixel, 16-bit depth, pixel size 0.63-micron, zoom 0.7). For the three-dimensional reconstruction, 40µm-thick z-stacks were acquired using 1µm-steps. Imaris-assisted reconstruction was performed as previously described (Althammer et al., 2020; Tang et al., 2020; Wahis et al., 2021). In brief, surface reconstructions were created based on the four individual channels (DAPI, OT, OTR-GFP, SYN/vGlut2). Co-localization of OT signal with SYN or vGlut2 was confirmed both manually and through the association/overlap function of IHC-labeled puncta in the Imaris software. IHC intensity of vGlut2 and SYN were assessed by creating spheres that precisely engulfed somata or dendrites as previously described (Tang et al., 2020).

### Ex vivo patch-clamp recording of vlPAG-OTR neurons

#### Slice preparation

To validate the functionality of vlPAG OTR neurons in OTR-iRES-Cre rats using electrophysiology, 12-13 week old female OTR-iRES-Cre rats (n = 4) received injections of the Cre dependent reporter virus, rAAV1/2-pEF1a-DIO-GFP, into the vlPAG. Following a 4-8 week recovery period, rats were anesthetized by administering i.p. ketamine (Imalgene 90 mg/kg) and xylazine (Rompun, 10 mg/kg). Transcardial perfusions were performed using an ice-cold, NMDG based aCSF was used containing (in mM): NMDG (93), KCl (2.5), NaH_2_PO_4_ (1.25), NaHCO_3_ (30), MgSO_4_ (10), CaCl_2_ (0.5), HEPES (20), D-Glucose (25), L-ascorbic acid (5), Thiourea (2), Sodium pyruvate (3), N-acetyl-L-cysteine (10) and Kynurenic acid (2). The pH was adjusted to 7.4 using either NaOH or HCl, after bubbling in a gas comprised of 95 % O_2_ and 5 % CO_2_. Rats were then decapitated, brains were removed and 350 µm thick coronal slices containing the hypothalamus were obtained using a Leica VT1000s vibratome. Slices were warmed for 10 minutes in 35°C NMDG aCSF then placed in a room temperature holding chamber filled with normal aCSF for at least 1 hour. Normal aCSF was composed of (in mM): NaCl (124), KCl (2.5), NaH_2_PO_4_ (1.25), NaHCO_3_ (26), MgSO_4_ (2), CaCl_2_ (2), D-Glucose (15), adjusted to pH 7.4 with HCL or NaOH and continuously bubbled in 95 % O_2_-5 % CO_2_ gas. Osmolarity of all aCSF solutions were maintained between 305-310 mOsm. Finally, slices were transferred from the holding chamber to an immersion-recording chamber and superfused at a rate of 2 ml/min.

#### Patch clamp recording

We targeted GFP+ or GFP-neurons in the vlPAG for whole-cell patch-clamp recording. The recording pipettes were visually guided by infrared oblique light video microscopy (DM-LFS; Leica). We used 4–9 MΩ borosilicate pipettes filled with a KMe based solution composed of (in mM): KMeSO_4_ (135), NaCl (8), HEPES (10), ATPNa_2_ (2), GTPNa (0.3). The pH was adjusted to 7.3 with KOH and osmolality was adjusted with sucrose to 300 mOsm/l, as needed. Data were acquired with an Axopatch 200B (Axon Instruments) amplifier and digitized with a Digidata 1440A (Molecular Devices, CA, USA). Series capacitances and resistances were compensated electronically. Data were sampled at 20 kHz and lowpass filtered at 5 kHz using the pClamp10 software (Axon Instruments). Further analysis was performed using Clampfit 10.7 (Molecular Devices; CA, USA) and Mini analysis 6 software (Synaptosoft, NJ, USA) in a semi-automated fashion (automatic detection of events with chosen parameters followed by a visual validation).

First spike latency quantification is measured as the duration preceding the first spike of a neuron submitted to a super-threshold stimulus of 50 pA.

#### TGOT stimulation

Finally, to validate the functionality of the putative OTR-Cre expressing cells in the vlPAG, we recorded GFP+ neurons in GAPfree (current clamp mode). Following a 5 min baseline recording period, a solution containing the OTR agonist, [Thr^4^Gly^7^]-oxytocin (TGOT, 0.4µM), was pumped into the bath for 20 s. The recording continued for a total of 20 min while the frequency of action potentials (APs) was quantified as the measure of neuronal activity. As a control, we repeated this procedure while patching GFP-neurons in the same vicinity. Neurons were held at ∼50 mV by injecting between 0 and 10 pA of current throughout the recording and all ex vivo experiments were conducted at room temperature.

### In vivo extracellular recording of vlPAG neurons

To test the effects of endogenous OT release on OTR cells of the vlPAG in vivo, female Wistar rats (n = 11) were injected with rAAV_1/2_-pOT-ChR2-mCherry into the PVN. After a 4-8 week recovery period, rats were anaesthetized with 4% isoflurane and placed in a stereotaxic frame before reducing the isoflurane level to 2%. A silicone tetrode coupled with an optical fiber (Neuronexus, USA) was inserted into the PAG to allow for stimulation of the ChR2 expressing axons of PVN-OT neurons projecting to vlPAG while recording the activity of putative OTR expressing neurons in the vicinity. Optical stimulation was delivered using a blue laser (λ 473 nm, output of 3 mW, Dream Lasers, Shanghai, China) for 20 s at 1 Hz, with a pulse width of 5 ms. Extracellular neuronal activity was recorded using a silicone tetrode coupled with an optic fiber (Q1x1-tet-10mm-121-OAQ4LP; Neuronexus, USA). Data were acquired on an MC Rack recording unit (Multi Channel Systems), and spikes were sorted by Wave Clus (Chaure et al., 2018). Spike data was analyzed with custom MATLAB (MathWorks) scripts and the MLIB toolbox (Stüttgen Maik, Matlab Central File Exchange).

The firing rate of each recording unit was smoothed by convolution of Gaussian distribution, whose width was 10 s and standard deviation was 5. The mean firing rate of the baseline period (BSmean) was defined as 0% activity and subtracted from the firing rates (FR) of the whole period (FR-BSmean). Maximum absolute activity (max(|FR-BSmean|)) was found using the highest absolute value among moving means of (FR-BSmean) with a time window of 21 s. When maximum absolute activity was found to exceed the BSmean, it was defined as 100% activity (cell#1∼21), whereas when maximum absolute activity was found to be below BSmean, it was defined as -100% activity (cell#22∼23).

### In vivo extracellular recording of WDR SC neurons

Adult female Wistar rats (n = 22) were anesthetized with 4% isoflurane and a then maintained at 2% after being placed in a stereotaxic frame. A laminectomy was performed to expose the L4-L5 SC segments, which were then fixed in place by two clamps positioned on the apophysis of the rostral and caudal intact vertebras. The dura matter was then removed. To record wide-dynamic-range neurons (WDR), a silicone tetrode (Q1x1-tet-5mm-121-Q4; Neuronexus, USA) was lowered into the medial part of the dorsal horn of the SC, at a depth of around 500 -1100 µm from the dorsal surface (see Figure 5A for localization of recorded WDRs). We recorded WDR neurons of lamina V as they received both noxious and non-noxious stimulus information from the ipsilateral hind paw.

We measured the action potentials of WDR neurons triggered by electrical stimulation of the hind paw. Such stimulation induced the activation of primary fibers, whose identities can be distinguished by their spike onset following each electrical stimuli (Aβ-fibers at 0-20 ms, Aδ-fibers at 20-90 ms, C-fibers at 90-300 ms and C-fiber post discharge at 300 to 800 ms). When the WDR peripheral tactile receptive fields are stimulated with an intensity corresponding to 3 times the C-fiber threshold (1 ms pulse duration, frequency 1 Hz), a short-term potentiation effect, known as wind-up (WU), occurs that leads to an increased firing rate of WDR neurons (Mendell and Wall, 1965; Schouenborg, 1984). Because the value of WU intensity was highly variable among recorded neurons within and across animals, we averaged the raster plots two dimensionally across neurons within each group of rats. We further normalized these data so that the plateau phase of the maximal WU effect was represented as 100 percent activity. As WU is dependent on C-fiber activation, it can be used as a tool to assess nociceptive information in the SC and, in our case, the anti-nociceptive properties of OT acting in the vlPAG. We recorded WDR neuronal activity using the following protocol: 40 s of hind paw electric stimulation to induce maximal WU followed by continued electrical stimulation to maintain WU while simultaneously delivering 20 s of vlPAG blue light stimulation (30 Hz, 10ms pulse width, output ∼3 mW), followed by another 230 s of electrical stimulation alone to observe the indirect effects of OT on WU in WDR neurons. Electrical stimulation was ceased after the 290 s recording session to allow the WU effect to dissipate. Following a 300 s period of no stimulation, the ability of the WU effect to recover was assessed by resuming electrical stimulation of the hind paw for 60 s of WU. After another 10 min period without stimulation, we sought to confirm the effects of vlPAG OT activity on WU intensity by injecting 600 nl of the OTR antagonist, dOVT, (d(CH2)5-Tyr(Me)-[Orn8]-vasotocine; 1µM, Bachem, Germany) into the vlPAG of the rats expressing ChR2, and repeated the protocol described above.

The spikes of each recording unit were collected as raster plots with the vertical axis showing the time relative to electric shock, and the horizontal axis showing the number of electric shocks. Next, the raster plots were smoothed by convolution of the Gaussian distribution (horizontal width = 100 ms, vertical width = 20 ms, standard deviation = 20. The total number of C fiber derived spikes occurring between 90 and 800 ms after each electric shock was counted. The spike counts were smoothed with a moving average window of 21 s and the window containing the maximal activity was defined as ‘100% activity’, which was then used to normalize the activity of each recording unit. Finally, the normalized percent activity from each recording unit was averaged for each experimental condition and plotted in figure 5.

### Determination of the phase of estrous cycle

To ensure consistency across studies (Eliava et. al., 2016), all electrophysiological recordings were conducted during the diestrus phase of the ovarian cycle, which we determined using vaginal smear cytology (Marcondes et al., 2002). Briefly, a micropipette filled with 100 µL of saline solution (NaCl 0.9%) was placed in the rats’ vagina and cells were dissociated by pipetting up and down at least three times. A drop of the smear was placed on a glass slide and observed using a light microscope with a 40x or 100x objective lens. Ovarian phase was determined based on the proportion of leukocytes, nucleated epithelial cells and anucleate cornified cells. Animals in metestus, proestrus and estrus phases were excluded from experiments and reintroduced once they reached diestrus.

### Behavioral tests

#### Optogenetics

For in vivo optogenetic behavioral experiments, we used a blue laser (λ 473nm, 100 mW/mm², DreamLasers, Shanghai, China) coupled to optical fiber patch cables (BFL37-200-CUSTOM, EndA=FC/PC, EndB=Ceramic Ferrule; ThorLabs, USA). Optical fiber probes (CFMC12L10, Thorlabs, USA) were bilaterally implanted into the vlPAG (Coordinates relative to bregma: ML= ±2.0mm, AP= -6.7mm, DV=-7.0mm, medio-lateral angle=10°) under isoflurane anesthesia (4% induction, 2% maintenance), and then stabilized with dental cement. Following a two-week recovery period, all animals received special handling to habituate them to the fiber connection routine. Optical stimulation to the vlPAG was delivered using a series of pulse trains (intensity = ∼3 mW, frequency = 30 Hz, pulse width = 10ms, duration = 20s) during all applicable behavioral experiments.

#### Chronic constriction of the sciatic nerve

To produce the model of chronic neuropathic pain, we surgically implanted a cuff around the sciatic nerve to induce a chronic constriction injury as previously described (Yalcin et al., 2014). Briefly, under isoflurane anesthesia (5%) via a facemask, an incision was made 3-4mm below the femur on the right hind limb and 10mm of the sciatic nerve was exposed. A sterile cuff (2 mm section of split PE-20 polyethylene tubing; 0.38 mm ID / 1.09 mm OD) was positioned and then closed around the sciatic nerve. The skin was then sutured shut to enclose the cuff and allow chronic constriction to occur.

#### Mechanical hyperalgesia

Mechanical sensitivity was assessed using calibrated digital forceps (Bioseb^®^, France) as previously described (Wahis et al., 2021). Briefly, the habituated rat is loosely restrained with a towel masking the eyes in order to limit environmental stressors. The tips of the forceps are placed at each side of the paw and gradual force is applied. The pressure required to produce a withdrawal of the paw was used as the nociceptive threshold value. This manipulation was performed three times for each hind paw and the values were averaged. After each trial, the device was cleaned with a disinfectant (Surfa’Safe, Anios laboratory^®^).

#### Inflammatory hyperalgesia

In order to induce peripheral inflammation, 100 μL of complete Freund adjuvant (CFA; Sigma, St. Louis, MO), was injected into the right hind paw of the rat. All CFA injections were performed under light isoflurane anesthesia (3 %). Edema was quantified by using a caliper to measure the width of the dorsoplantar aspect of the hind paw before and after the injection of CFA. In effort to reduce the number of animals used, we did not include an NaCl-injected group, as it has already been shown that the contralateral hind paw sensitivity is not altered by CFA injection (Hilfiger et al., 2020).

#### Thermal hyperalgesia

To test the animal thermal pain sensitivity threshold, we used the Plantar test with the Hargreaves method (Ugo Basile^®^, Comerio, Italy) to compare the response of each hind paw of animals having received unilateral intraplantar CFA injection. The habituated rat is placed in a small box and we wait until the animal is calmed. We then exposed the hind paw to radiant heat and the latency time of paw withdrawal was measured. This manipulation was performed three times for each hind paw and the values were averaged. After each trial, the device was cleaned with a disinfectant (Surfa’Safe, Anios laboratory).

#### Conditioned Place Preference

The device is composed of two opaque conditioning boxes (rats: 30x32 cm; mice: 22x22 cm) and one clear neutral box (30x20 cm). Animals were tracked using a video-tracking system (Anymaze, Stoelting Europe, Ireland) and apparatus was cleaned with disinfectant (Surfa’Safe, Anios laboratory) after each trial. The animals underwent CPP as previously described (Wahis et al., 2021). Briefly, all rats underwent a three-day habituation period during which they were able to freely explore the entire apparatus for 30 min. On the 3^rd^ habituation day, exploration behavior was recorded for 15 min to determine the animals’ innate side preference. On the 4^th^ day, animals were injected with saline (i.p, 1 mL/kg) and placed in their innately preferred chamber (unpaired box) for 15 min. Four hours later, animals were injected with DCZ (i.p, 100 µg/kg at 1 mL/kg (Nagai et al., 2020)) to stimulate vlPAG OTR neurons expressing hM3D(Gq), and then placed in the innately non-preferred chamber (paired box) for 15 min. On the 5^th^ day, the animals were placed in the neutral chamber and allowed to explore the entire apparatus for 15 min. To control for potential locomotor effects, the total distance travelled during the test period was quantified and compared between all groups (Figure 6I).

### Drugs

All drugs used in this study are listed in the reagents and resource table.

### Statistical Analysis

For ex vivo electrophysiology data, one-way analysis of variance (ANOVA, Friedman test) followed by Dunn’s post-hoc multiple comparisons test to compare averages across the three conditions (Baseline vs TGOT vs Wash). Differences were considered significant at p<0.05. For in vivo electrophysiology data analysis, a paired-sample t-test was used to compare the average spike rates between the baseline and peak activity of PAG neurons in response to BL stimulation (**Figure 4**). A nonparametric, unpaired Wilcoxon rank sum test was used to compare the reduction in discharge of SC neurons between the wild type and the ChR2-expressing animals (**Figure 5**). A paired Wilcoxon signed rank test was used to compare the reduction discharge of SC neurons in the ChR2-expressing animals, between the “without dOVT” and “with dOVT” conditions (**Figure 5**). Wilcoxon rank sum tests were used to compare the latencies to reach the maximum, minimum, and the half value between the wild type and the ChR2-expressing rats (**Figure S5**).

Behavioral data are expressed as mean ± standard deviation (SD). Statistical tests were performed with GraphPad Prism 7.05 (GraphPad Software, San Diego, California, USA). All behavioral data failed the Shapiro-Wilk normality test. A Kruskall Wallis test with Dunn’s post-hoc test was used to compare the percent co-localization of c-Fos+ and GFP+ cells across pain conditions (Figure 6A). A repeated-measures non parametric mixed model ANOVA was used to analyze the effect of DCZ on pain in PAG OTr/DREADDGq neurons with Dunnett’s post-hoc multiple comparisons test to compare the effect of the (treatment) before (t = 0) and after the i.p. injection (20, 40, 60 and 180 min) (**Figure 6, S6**).

The ΔCPP score was calculated with the following formula in order to control for time spent in the neutral chamber: ΔCPP score = (paired_postcond_ - unpaired_postcond_) - (paired_hab_ - unpaired_hab_). A Mann Whitney test was used to compare the effect of DCZ on CPP difference score across the two groups as well as the distance travelled by the two groups (**Figure 6**). Differences were considered significant at p<0.05. Asterisks are used to indicate the significance level: * 0.01 ≤ p < 0.05, ** 0.001 ≤ p < 0.01, *** p < 0.001. All rats with off-target viral injection sites were removed from analysis.

**Figure S1 (related to Figure 1).**

**(A)** Detailed schema of the generation of knock-in OTR-Cre rats, displaying the genomic structure of the OTR gene, the targeting vector, and the insertion site.

**(B)** Injection schema for the labelling of OTR neurons in the generated OTR-Cre rats. The rAAV-pEf1a-DIO-GFP is injected into the PAG, labelling exclusively OTR-neurons.

**(C)** Images showing GFP (green) and DAPI (blue) staining, at various distances from Bregma, in the vlPAG of OTR-Cre rats injected with the Ef1a-DIO-GFP virus. Scale bar = 100µm. Aq = Aqueduct.

**(D)** TGOT-induced increase of AP frequency of GFP-negative neurons in the PAG. **D1** Example traces from a GFP-negative cell under baseline, TGOT application, and wash out conditions. **D2** Quantification of TGOT effect on AP frequency of GFP-negative neurons (Baseline 0.119 ± 0.046 Hz vs TGOT 0.108 ± 0.049 Hz vs Wash 0.122 ± 0.064 Hz; Friedman’s test = 1.6, p = 0.48, n = 9). Data are represented as mean ± sem and as individual paired points.

**(E)** TGOT-induced change in first spike latency (FSL) of GFP-negative neurons in the PAG. **E1**. FSL quantification for GFP-negative neurons (OTR^-^): no differences were found between the baseline condition (31.95 ± 9.44 ms) and after the TGOT incubation 37.08 ± 14.89 ms, non significative (n.s.) p = 0.788 (paired two-tailed t test, n = 7). Data are represented as mean ± SEM and as individual paired points. **E2**. Proportion of neuron after TGOT incubation with a decrease of the FSL (n = 1/7, blue); an increase (n= 2/7, red) or without a variation of ± 10 ms (no effect, gray (n = 4/7). **E3**. Representative evoked currents in a GFP-negative neuron in response to a square current step (50 pA) in baseline (black line) and oxytocin (blue line) conditions

**Figure S2 (related to Figure 2).**

**(A)** Distinction of parvocellular (parvOT) and magnocellular (magnOT) OT-neuron populations. **A1** Schema for the identification of magnOT and parvOT neurons: injection of green retrobeads into the PAG, staining for OT (red) and central application of fluorogold (FG, blue), which labels only magnOT neurons. **A2** Image of retrolabeled neurons containing fluorescent beads (green) as well as staining for Fluorogold (blue) and OT (red). White arrows indicate the co-localization of OT and FG (magnOT neuron) and the white asterisk indicates the co-localization of retrobeads and OT. Scale bars = 25 µm. n = 3 female rats.

**(B)** Retrograde labelling of projections from PVN OT-neurons to the vlPAG and SC. **B1** Schema of the experimental design with injection site of green retrobeads in the vlPAG (**B2**) and red retrobeads in the SC (L5, **B3**). Scale bars = 200 µm. **B4** Image of PVN OT-neurons (blue) projecting either to the vlPAG (green, white asterisk) or to the SC (red, white arrow). No OT neuron containing both red and green retrobeads was found. Aq = Aqueduct. 3v = third ventricle. Scale bar = 50µm. n = 4 female rats.

**(C)** PAG-projecting PVN OT-neurons from Figure 2B do not target the SC or SON. **C1** Schema of viral injection showing injection of rAAV-pOT-DIO-GFP in the PVN and rAAV-CAV2-Cre in the vlPAG of WT rats. **C2** Images of slices from the spinal cord at different levels (cervical, C3; thoracic, T7; lumbar, L4) and from the SON. No GFP fibers were found. Scale bars = 200 µm. n = 2 female rats.

**(D)** SON-projecting PVN OT-neurons target the SC but not the vlPAG. **D1** Schema of viral injection showing injection of rAAV-pOT-DIO-GFP in the PVN and rAAV-CAV2-Cre in the SON of WT rats. **D2 Left** Images of PVN OT-neurons (red) expressing GFP (green) projecting fibers to the SC (L5, NeuN in red, scale bars = 100 µm). **Right** Image of the vlPAG showing no GFP positive fibers. Scale bar = 100 µm. n = 4 female rats.

**(E)** SC-projecting PVN OT-neurons target the SON but not the vlPAG. **E1** Schema of viral injection showing injection of rAAV-pOT-DIO-GFP in the PVN and rAAV-CAV2-Cre in the SC of WT rats. **E2 Left** Images of PVN (scale bars = 40 µm) OT-neurons (red) expressing GFP (green) projecting fibers to the SON (scale bar = 100 µm). opt = optic tract. **Right** Image of the vlPAG showing no GFP positive fibers. Aq = aqueduct. Scale bar = 100 µm. n = 4 female rats.

**Figure S3 (related to Figure 3).**

**(A)** Pipeline for Imaris-assisted 3D reconstruction based on raw IHC images.

**(B)** Confocal images showing OT fibers and OTR-positive neurons in the vlPAG. **B1** Images of OTR neurons (green), OT fibers (magenta), vGluT2 (red) and DAPI (blue). Scale bar = 100 µm. **B2** Overlay. Scale bar = 100 µm.

**(C)** Automated quantification of the number of SYN positive voxels in GFP-positive and negative neurons reveals no difference in overall synaptic input. n = 8 images per animal, p=0.6689.

**(D)** Analysis of somatic volume reveals no difference between OTR-positive (n = 496) and OTR-negative (n = 3840) soma sizes. n = 8 images per animal, p = 0.8104

**Figure S4 (related to Figure 4).**

**(A)** Visualization of the last step of spike sorting-superparamagnetic clustering (ascription of spikes to their corresponding neurons). This step is preceded by 1) automatic spike detection based on amplitude threshold and feature extraction – wavelets coefficients. From left to right: superimposed clusters, cluster separation and unassigned spikes based on spike shapes

**(B)** Spike shapes represented in each channel of the tetrode.

**(C)** Cluster 3 (from B) detailed profile, depicting (from left to right, and up to down) spikes’ waveforms, baseline & peak voltages, spikes waveform variability over time, spikes parameters, stability presented as spikes count/time, activity during interspikes intervals of different length, density plots, waveform of selected spikes, PSTH, waveforms detected around an event and spikes distribution

**(D)** Onset, peak and offset of BL-induced excitation in the vlPAG. Onset is defined as the first time point when the spike rate exceeded the threshold (BS_mean_ + 4BS_SD_); BS_mean_ represents the mean firing rate of the baseline period, and BS_SE_ is the standard deviation of firing rate during the baseline period. Offset is defined as the last time point when the spike rate fell below the threshold for more than 20 s during the 300 s following BL (n = 21).

**(E)** Number of active cells in each single second. “Active” defined by a spike rate above the threshold (BS_mean_ + 4BS_SD_) (n = 21).

**(F)** Image showing a representative recording site in the vlPAG. Dotted line indicates placement of the optic fiber. Aq = aqueduct. Scale bar = 100 µm.

**Figure S5 (related to Figure 5).**

Box plot showing maximum activity, minimum activity, and half activity latencies. Whiskers indicate the minimum and maximum latency. Colored box shows the range between 1st and 3rd quantile, median is depicted as vertical line. [(1^st^ quartile, median, 3^rd^ quantile); WT:(26.00, 31.50, 46.00), ChR2: (26.25, 35.00, 55.75)]; latency to reach the half reduction of WU (s) [WT: (75.50, 96.50, 163.00), ChR2 (64.25, 83.00, 125.75)]; or latency to reach the maximum reduction of WU (s) [WT (183.00, 209.00, 263.00), ChR2 (198.50, 250, 269.75)].

**Figure S6 (related to Figure 6).**

**(A)** Optogenetic stimulation of the vlPAG in the chronic constriction of the sciatic nerve (CCI) model of neuropathic pain. **A1** Schema of the injection of rAAV-pOT-ChR2-mCherry in the PVN and optic fiber implantation in the PAG. **A2-3** Graphs showing the threshold response to mechanical stimuli in female rats optogenetically stimulated in vlPAG (as in Figure 6C and 6D). The effect of vlPAG-BL was measured at 5 min, 1 h and 3 h after vlPAG-BL on the CCI side (**A2**) as well as contralateral side (**A3**). All results are expressed as average ± SEM. * p < 0.01, ** p < 0.01, n = 7.

**(B)** Verification for the expression of Gq-mCherry in the vlPAG. **B1** Schema of the injection of rAAV-pEf1a-DIO-Gq-mCherry in the PAG. **B2** Image showing the expression of rAAV-Ef1A-DIO-Gq-mCherry (red) with staining for DAPI (blue). Scale bar = 100 µm. Aq = aqueduct.

**(C)** Mechanical pain threshold after DCZ administration in the CFA (**C1**) and contralateral (**C2**) paw of male rats expressing Gq-mCherry (blue) or mCherry only (gray) in vlPAG OTR neurons. All results are expressed as average ± SEM. *** p < 0.0001, n = 6 per group.

**(D)** Thermal pain threshold after DCZ administration in the CFA (**D1**) and contralateral (**D2**) paw of male rats expressing Gq-mCherry (red) or mCherry only (gray) in vlPAG OTR neurons. All results are expressed as average ± SEM. *** p < 0.0001, n = 6 per group.

**(E)** Schema depicting the experimental timeline of the CPP protocol.

| REAGENT OR RESOURCE | SOURCE | IDENTIFIER |
| --- | --- | --- |
| **Antibodies** |  |  |
| Chicken anti-GFP primary antibody | Abcam | ab13970 |
| Mouse NeuN primary antibody | Millipore | MAB377 |
| Rabbit vGluT2 primary antibody | Synaptic systems | 135 103 |
| Rabbit anti-dsRed primary antibody | Living Colors | 632496 |
| GAD67 | Millipore | MAB5406 |
| Guinea-pig anti-Fluorogold primary antibody | Protos Biotech Corp | NM-101 |
| Mouse c-fos polyclonal primary antibody | Santa-Cruz | sc-8047 |
| Mouse monoclonal anti-OT primary antibody | Provided by Dr. Harold Gainer | PS 38 |
| DAPI | Vector Laboratories | H-1200-10 |
| Synaptophysin | Abcam | Ab32127 |
| Rabbit polyclonal anti-vGluT2 primary antibody | SYSY | 135403 |
| **Bacterial and Virus strains** |  |  |
| rAAV1/2-OTp-Venus | Lab made | N/A |
| rAAV1/2-OTp-mCherry | Lab made | N/A |
| rAAV1/2-OTp-FLEX-GFP | Lab made | N/A |
| CAV2-CMVp-Cre | Montpellier Vectorology Platform | N/A |
| rAAV1/2-OTp-ChR2-mCherry | Lab made | N/A |
| rAAV1/2-Ef1ap-FLEX-GFP | Lab made | N/A |
| rAAV1/2-Ef1ap-FLEX-mCherry | Lab made | N/A |
| rAAV1/2-Ef1ap-FLEX-hM3D(Gq)-mCherry | Lab made | N/A |
| **Chemicals, Peptides, and Recombinant Proteins** |  |  |
| Fluorogold™ | Santa-Cruz | sc-358883 |
| Oxytocin Receptor Antagonist | Merck | L-368,899 |
| Deschloroclozapine | MedChemExpress | 1977-07-7 |
| TGOT | Bachem® | 50-260-164 |
| CFA | Sigma | 32160405 |
| dOVT | Bachem | d(CH2)5-Tyr(Me)-[Orn8]-vasotocine |
| Retrobeads™ | Lumaflour | N/A |
| **Experimental Models: Organisms/Strains** |  |  |
| Rattus Norvegicus (Wistar) | Janvier | N/A |
| Rattus Norvegicus (Sprague Dawleys) | Charles River | N/A |
| OTR-IRES-Cre (Sprague Dawleys) | Lab made | N/A |
| **Software and Algorithms** |  |  |
| Graphpad prism 7.05 | www.graphpad.com | N/A |
| Fiji | www.imagej.net/Fiji | N/A |
| Adobe Photoshop CS5 | www.adobe.com | N/A |
| Adobe Illustrator 16.05 | www.adobe.com | N/A |
| Matlab | Mathworks | N/A |
| **Other** |  |  |
| Optic fiber implants | www.thorlabs.com | CFMC12L10 |
| 473nm Blue Laser Generator | www.dreamlasers.com | SDL-473-XXXT |
| Programmable Pulse Stimulator (A.M.P.I.) | www.ampi.co.il | Master-9 |

