## Supplementary figures and images for "A novel analgesic pathway from parvocellular oxytocin neurons to the periaqueductal gray"

### Supplementary Figures

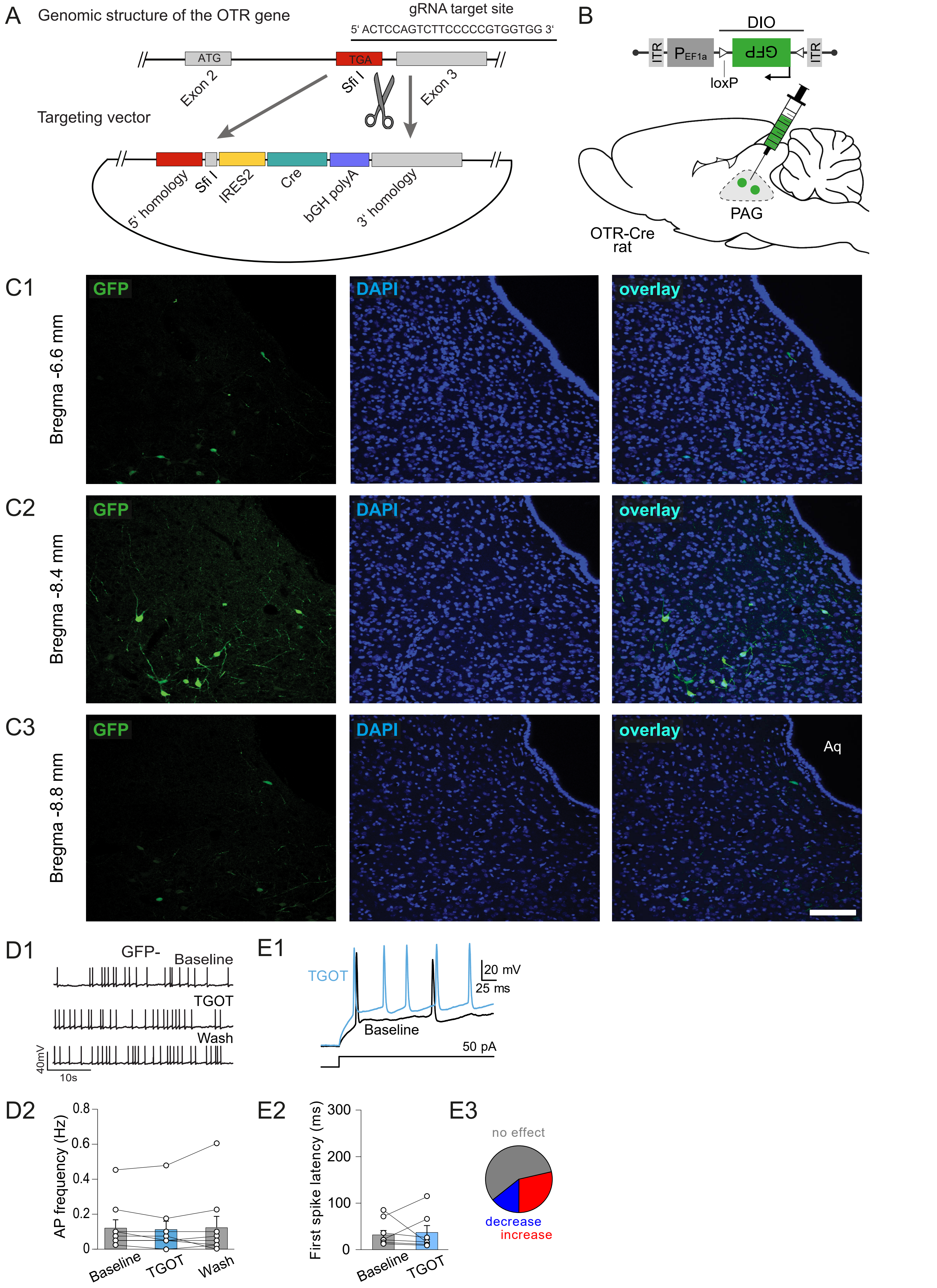

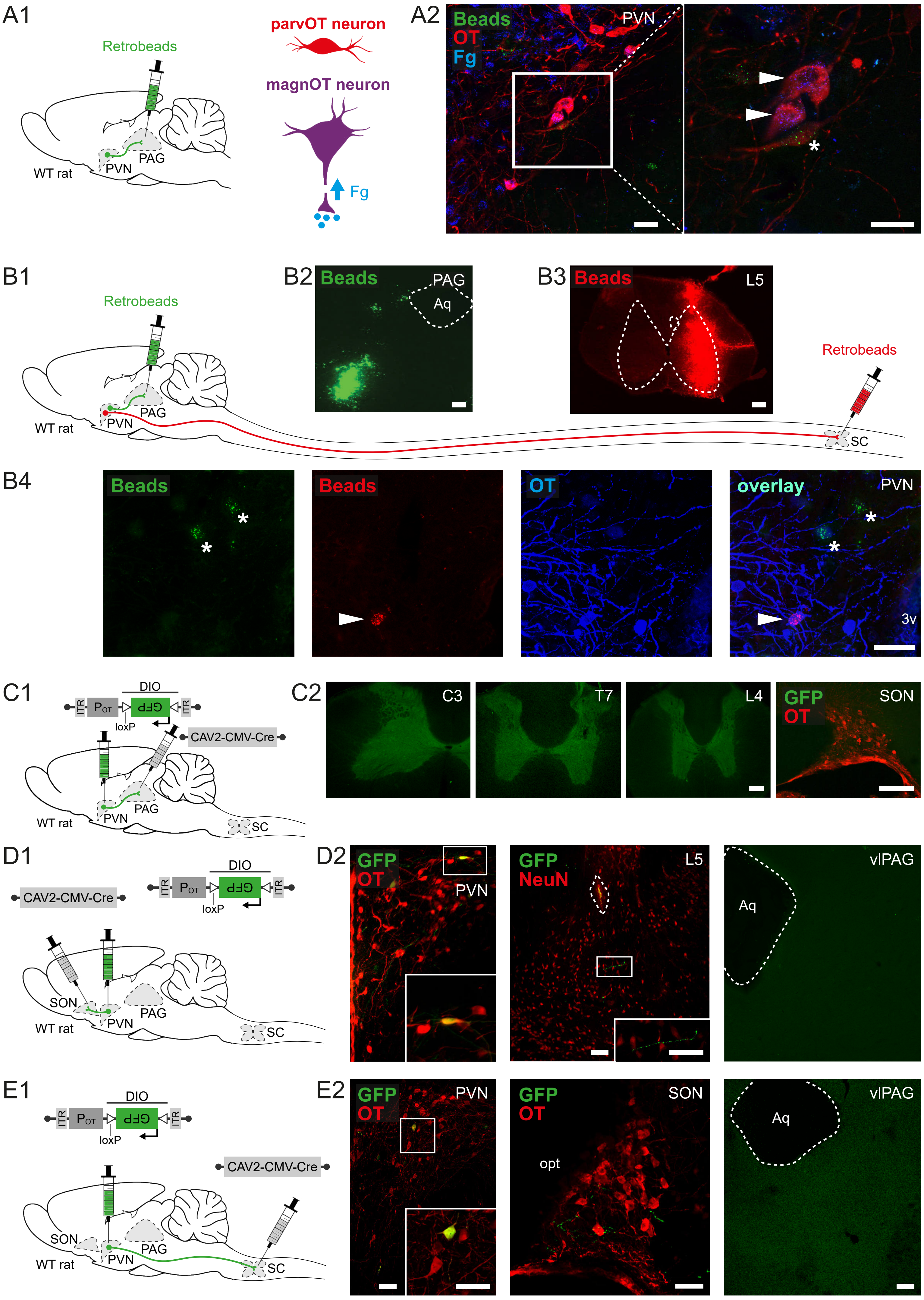

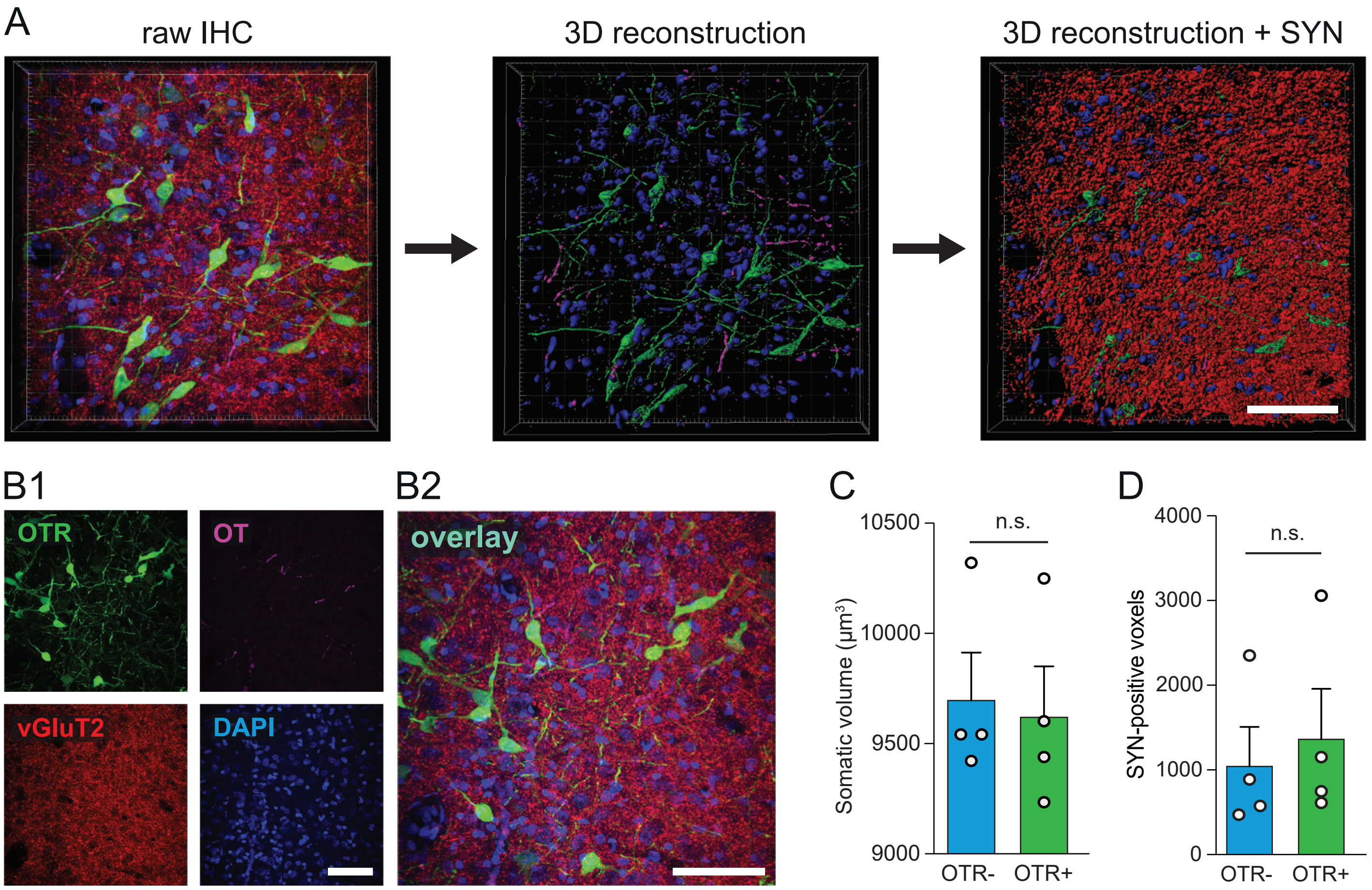

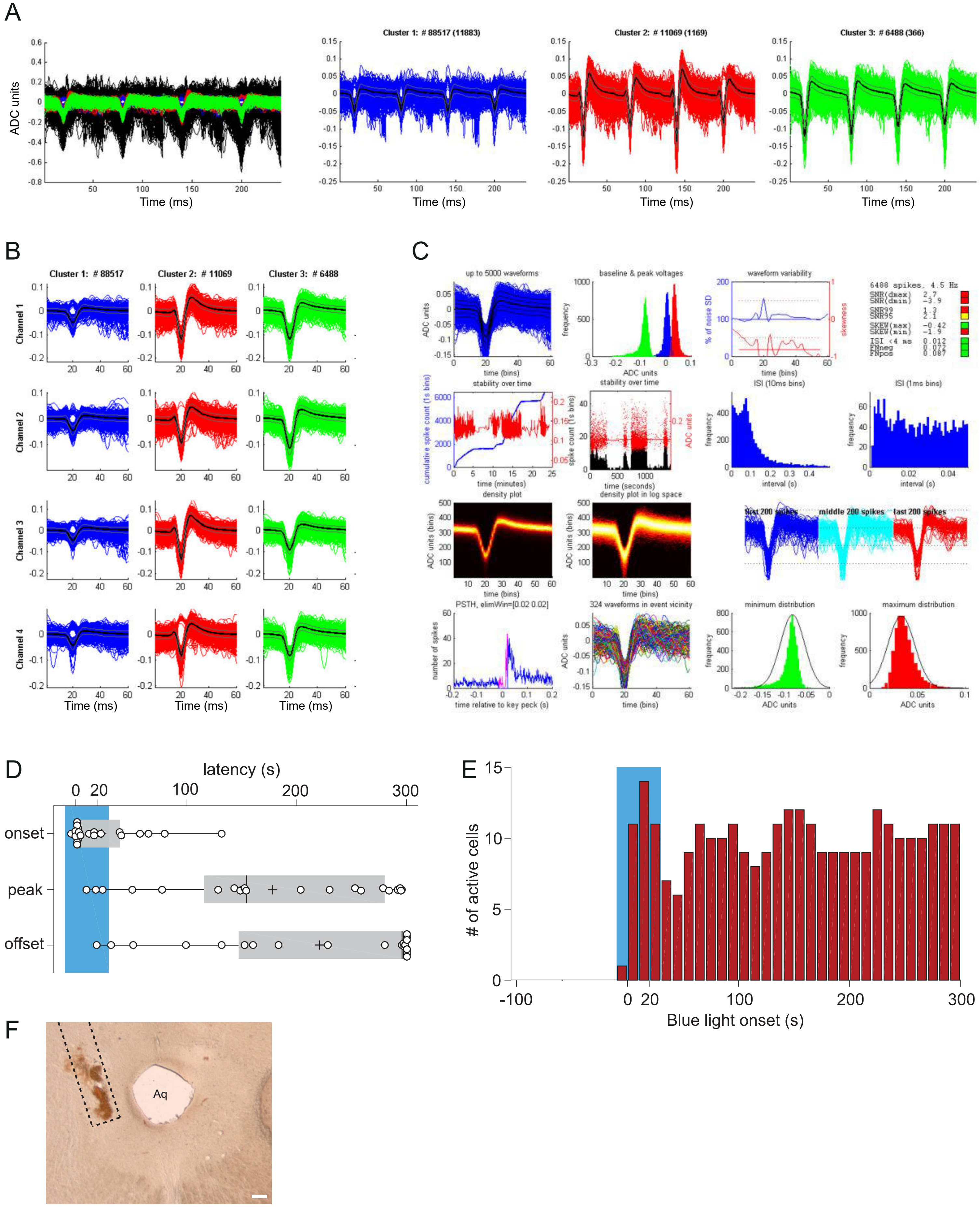

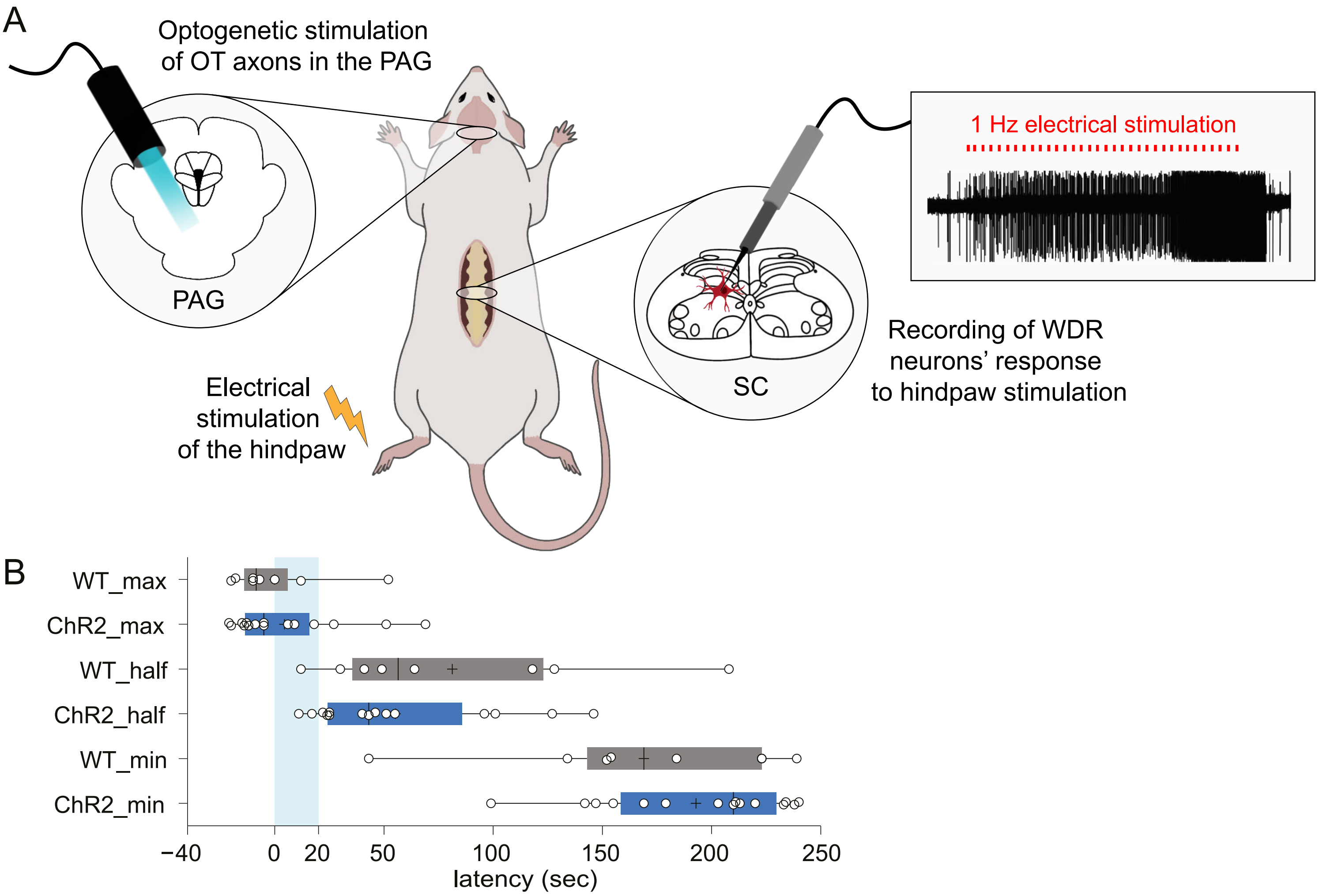

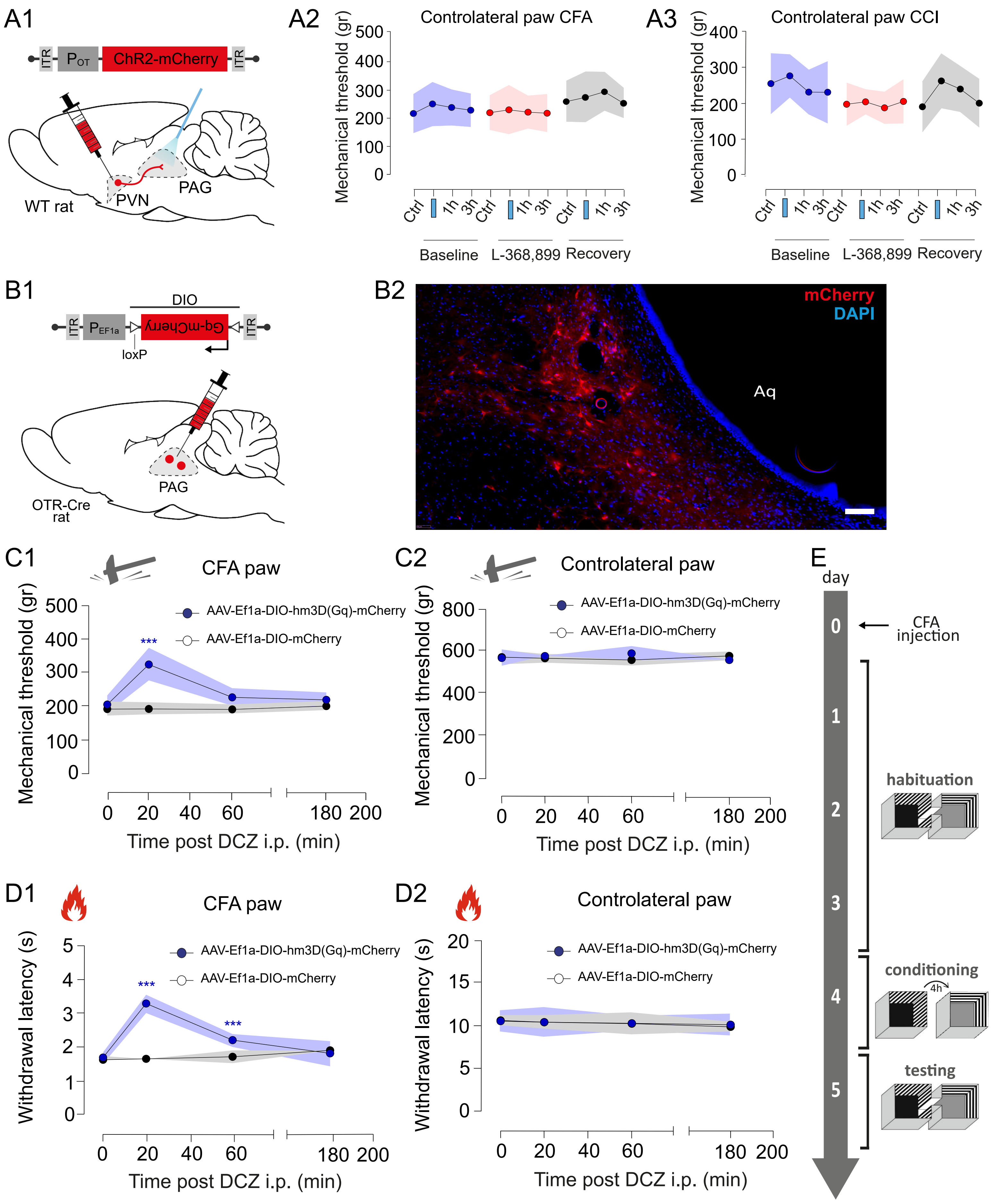
